# Early mitophagy defects and impaired mitochondrial energy metabolism drive target organ damage progression: lessons from the Fabry heart

**DOI:** 10.64898/2026.04.15.718770

**Authors:** Jessica Gambardella, Antonella Fiordelisi, Federica Andrea Cerasuolo, Antonietta Buonaiuto, Roberta Avvisato, Alessandro Viti, Eduardo Sommella, Pietro Campiglia, Valeria D’Argenio, Nella Prevete, Antonio Pezone, Stefania D’Apice, Giovanna Giuseppina Altobelli, Fahimeh Varzideh, Shivangi Pande, Roberta Paolillo, Cinzia Perrino, Eleonora Riccio, Antonio Pisani, Antonio Bianco, Junichi Sadoshima, Letizia Spinelli, Gaetano Santulli, Daniela Sorriento, Guido Iaccarino

## Abstract

Increased literature support the pathogenetic role of dysfunctional energetic metabolism in the setup and progression of organ damage and failure. Genetic diseases often offer the possibility to investigate pathogenetic mechanisms. In particular, excessive cardiac damage is the most frequent cause of mortality in Fabry disease (FD), a genetic condition caused by deficient α-galactosidase A (GLA) activity, leading to globotriaosylceramide (Gb3) accumulation. Beyond Gb3 storage, metabolic alterations and mitochondrial dysfunction, supported by in vitro evidence or studies in other tissues, may contribute to FD cardiomyopathy. This study investigated, for the first time, the mechanisms of mitochondrial involvement in FD, its role in determining cardiac manifestations, and its potential as a therapeutic target. We used a humanized FD mouse model (R301Q-Tg/GLA knockout), along with derived embryonic fibroblasts and neonatal and adult cardiomyocytes, to assess mitochondrial function across the lifespan. FD cells showed impaired mitophagy, reduced mitochondrial respiration, and increased reactive oxygen species production. Importantly, this mitochondrial dysfunction exacerbated the lysosomal deficit in FD cells, forming a vicious cycle. In cardiomyocytes, these alterations progressed with age, leading to the accumulation of dysfunctional mitochondria, energetic failure, and, in adult hearts, terminal mitochondrial damage and apoptosis. These events ultimately result in cardiac remodeling and dysfunction, including hypertrophy and diastolic impairment. Indeed, L-arginine supplementation, which promotes NO/PGC-1α–dependent mitochondrial rescue, prevented the development of cardiac abnormalities in FD mice. Our findings identify early mitochondrial dysfunction as a key driver of FD cardiomyopathy and support mitochondrial targeting, including L-arginine supplementation, as a promising adjuvant therapeutic strategy. The mechanistic link between lysosomal dysfunction, altered mitochondrial turnover, and energetic collapse emerges as a key targetable pathway in organ damage, extending beyond FD.

**Graphical abstract:** 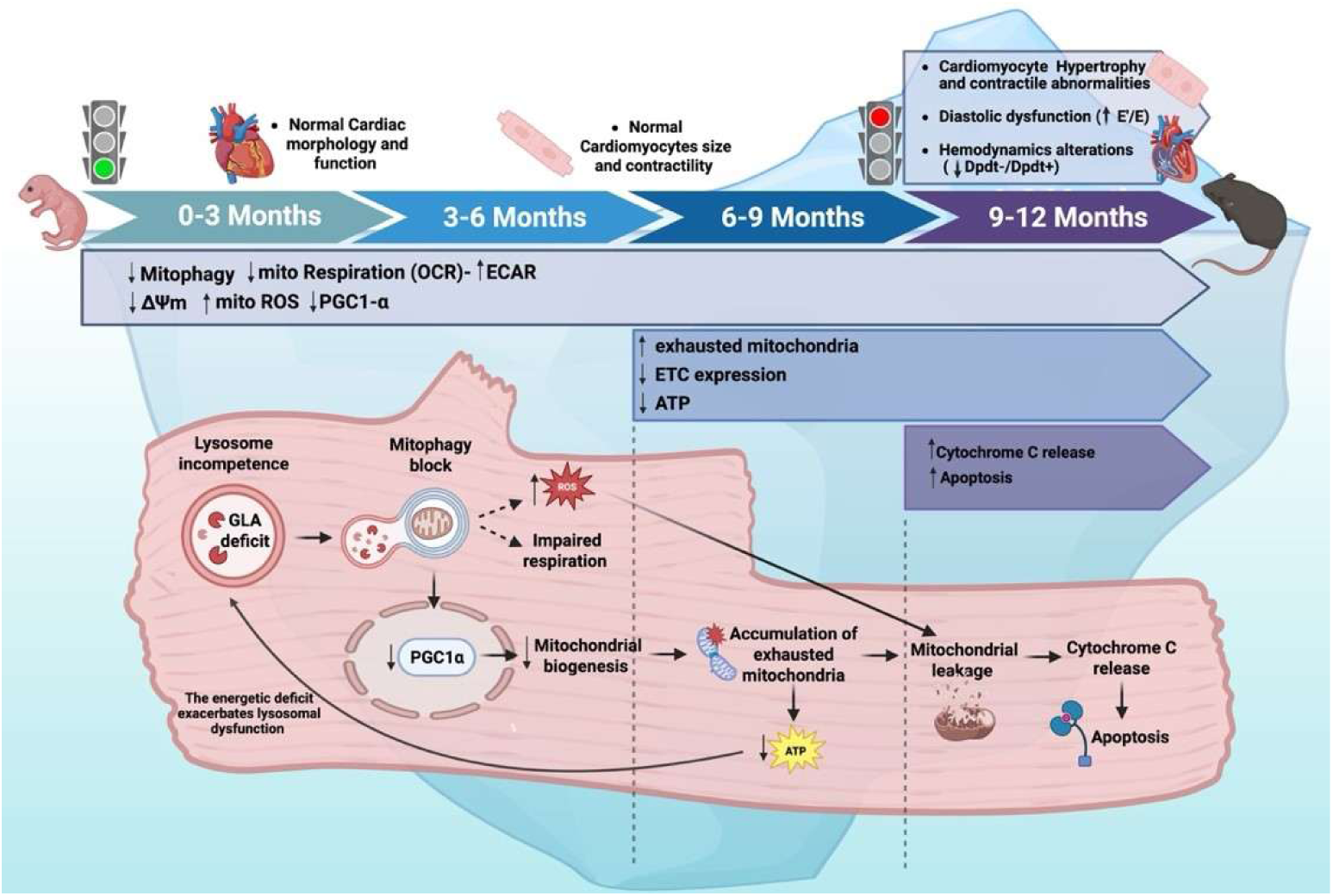

**Cardiac manifestations vs mitochondrial alterations in Fabry disease: the visible tip and the hidden base of the iceberg:** Cardiac manifestations in hR301Q Tg/KO mice become evident from 9 months of age. However, mitochondrial homeostasis is perturbed much earlier (neonatal to young stages), with impaired mitophagy, reduced mitochondrial respiration and membrane potential, increased ROS production and PGC-1α downregulation. At later stages, from 6 months of age, mitochondrial dysfunction progresses and begins to impact cellular energetics, as indicated by reduced ETC expression and the onset of energetic deficit (ATP reduction). The resulting energetic collapse, together with progressive mitochondrial leakage, leads to cardiomyocyte hypertrophy, apoptosis, and dysfunction, which become detectable from 9 months of age, when clinical signs emerge. These findings support a mechanistic model in which 1) lysosomal incompetence due to GLA deficit is the initiating event inducing impairment of mitophagy; 2) Unsuccessful mitophagy, induces downregulation of PGC-1a-dependent mitogenesis; 3) exhausted mitochondria accumulate, inducing energetic collapse (able to exacerbate lysosomal dysfunction and further perturb mitophagy in a vitious cycle); 4) ultimate mitochondrial leakage induces Cytochrome C release and apoptosis activation. This cascade of molecular events is responsible for clinical manifestations, and mitochondrial targeting prevents cardiac organ damage.

**Significance statement:** Fabry disease is a rare genetic disorder in which cardiac complications are a major cause of death, yet underlying mechanisms remain unclear. Here, we identify mitochondrial dysfunction as an early pathogenic event associated with impaired mitophagy, whereby defective mitochondrial quality control both results from and exacerbates lysosomal dysfunction, creating a self-reinforcing cycle that drives disease progression. Using a humanized model, we demonstrate that mitochondrial dysfunction is a key determinant of cardiac phenotype in vivo, driving energetic failure, oxidative stress, and cardiac damage. Importantly, L-arginine treatment restores mitochondrial function and prevents cardiac abnormalities. Our findings define a broadly relevant pathogenic axis linking lysosomal dysfunction, mitophagy failure, and mitochondrial impairment, that lead to impaired energetic metabolism and consequent cardiac hypertrophy, independently from GB3 accumulation. The implications of our study go beyond Fabry disease and support the therapeutic targeting of cellular energy homeostasis to prevent and treat organ damage and failure in chronic diseases.

**IMPORTANT:**

- Manuscripts submitted to Review Commons are peer reviewed in a journal-agnostic way.
- Upon transfer of the peer reviewed preprint to a journal, the referee reports will be available in full to the handling editor.
- The identity of the referees will NOT be communicated to the authors unless the reviewers choose to sign their report.
- The identity of the referee will be confidentially disclosed to any affiliate journals to which the manuscript is transferred.

**GUIDELINES:**

- For reviewers: https://www.reviewcommons.org/reviewers
- For authors: https://www.reviewcommons.org/authors

**CONTACT:** The Review Commons office can be contacted directly at

## Introduction

Fabry disease (FD) is the most prevalent lysosomal storage disorder (LSD), caused by genetic deficiency of a-Galactosidase-A (GLA) [1]. This enzymatic deficit disrupts glycosphingolipid metabolism and leads to cellular accumulation of globotriaosylceramide (Gb3) across multiple organs. FD reduces life expectancy, and its clinical manifestations range from early cardiac and multiorgan dysfunction to later-onset forms predominantly affecting the heart [2, 3]. Enzyme replacement therapy (ERT) is the current standard treatment; however, it has significant adverse effects and limited efficacy in managing FD-related cardiomyopathy, which remains the leading cause of death [4, 5] [6, 7]. Cardiac manifestations are heterogeneous, ranging from diastolic and systolic dysfunction to conduction abnormalities, myocardial fibrosis, valvular disease, and left ventricular hypertrophy [8]. Notably, diastolic dysfunction can occur early in the disease, preceding other cardiac signs.

The exact pathogenesis of FD cardiomyopathy has not been fully elucidated. Indeed, Gb3 deposits account for only 1–2% of total cardiac mass, supporting the hypothesis that, beyond mechanical storage, cardiac involvement results from activation of multiple signaling pathways [9]. The identification of these early mechanisms is crucial to address reversible pathological changes. Among the candidate pathogenic mechanisms, mitochondrial dysfunction has recently hypothesized as a potential contributor to FD pathophysiology [10], based on evidence of mitochondrial alterations reported in Fabry disease. Indeed, defects in mitochondrial function—specifically in oxygen consumption rate and electron transport chain (ETC) activity—have been described in PBMCs and fibroblasts from Fabry patients, along with reduced oxidative metabolism in skeletal muscle from preclinical models of **FD** [11, 12]. However, to date, studies investigating mitochondrial dysfunction in FD remain largely descriptive, as they have not addressed the underlying mechanisms, its pathogenic role in driving organ damage in vivo at the pathophysiological level, or its potential as a therapeutic target. Considering the central role of mitochondria in myocardial homeostasis, these knowledge gaps are particularly critical in the Fabry heart, where disease mechanisms remain poorly understood and current therapies are only partially effective.

Mitochondria and lysosomes are mutually associated [13]. In particular, the lysosome is pivotal for exhausted-mitochondria scavenging, and mitochondria regeneration [11]. We hypothesize that defective lysosomal function in FD leads to early impairment of mitochondrial turnover, resulting in the accumulation of exhausted mitochondria. This, in turn, promotes energetic collapse and oxidative stress, triggering maladaptive responses in tissues with high energetic demand and mitochondrial-dependent metabolism, including the heart, and ultimately driving the clinical manifestations of the disease.

Here, we verify this hypothesis using a murine model of FD, the hR301Q Tg/KO mouse (FD mouse) [14]. This mouse lacks murine *GLA* and expresses the R301Q variant of the human GLA gene causative of classic and late-onset FD. The hR301Q Tg/KO model effectively reproduces the accumulation of GB3 in disease-relevant tissues, including the heart. For mechanistic studies ascertaining alterations of mitochondrial function and turnover, we used mouse embryonic fibroblasts (MEFs) isolated from FD and WT mice, an exemplificative and manipulable *in-vitro* model. Then, we assessed the onset of cardiac manifestations in FD mice *in vivo* and *ex vivo*, exploring the association with cardiac energetics and mitochondrial alterations. Finally, we evaluated whether mitochondrial targeting can prevent FD cardiac manifestations using L-Arginine (Arg) supplementation, known to improve mitochondrial function, turnover, and energetics, especially in myopathy and cardiometabolic disorders [15, 16]. Importantly, we provide proof of concept that mitochondrial dysfunction exacerbates lysosomal impairment in FD, emerging as an early and primary contributor to disease pathogenesis.

Overall, our findings demonstrate that mitochondrial dysfunction is not merely a secondary consequence of cellular damage in FD, but an early and active driver of organ injury, and a targetable process to prevent clinical manifestations. These results support a paradigm shift in the management and treatment of FD, and more broadly define a causal and mechanistic axis linking lysosomal dysfunction, impaired mitophagy, mitochondrial quality control, energetic failure, and organ damage—an axis with relevance well beyond Fabry disease.

## Results

### Mitochondrial dysfunction and impaired mitochondrial quality control in FD cells

In order to evaluate MEFs as a potential cellular model for studying mitochondrial biology impacted by impaired GLA activity, we conducted a comprehensive series of imaging, functional, and molecular approaches. Basal and maximal oxygen consumption rate (OCR), assessed by mitostress test, were significantly impaired in FD MEFs compared to WT cells. This observation was corroborated by the evaluation of mitochondrial energetic reserve (spare capacity) and ATPlinked respiration, providing compelling evidence of mitochondrial dysfunction (**Fig. 1A**). The increase of Extracellular Acidification Rate (ECAR) in FD MEFs, suggested a compensatory activation of glycolysis in response to the defective oxidative phosphorylation (OXPHOS) **(Fig.S1 A).** Accordingly, cellular ATP content was significantly reduced in FD MEFs (**Fig.1 B**), while the production of mitochondrial Reactive Oxygen Species (ROS) was markedly elevated (**Fig. 1C**). Additionally, Cytochrome C release in the cytosol, a known marker of mitochondrial permeabilization and leakage, was augmented in FD MEFs (**Fig. 1D**). Confocal microscopy revealed a reduction of elongated mitochondria and an increased pool of fragmented mitochondria (high circularity), devoid of functional structure and interconnections (**Fig. 1E**). We also observed significant alterations of mitochondrial membrane potential (TMRE staining) compatible with the reduced mitochondrial oxidative phosphorylation capacity and ROS overproduction (**Fig. 1F**). The mitochondrial quality control (mQC) is a process that maintains healthy mitochondria by: i) fission (fragmentation) of damaged mitochondrial portions, ii) their elimination by lysosomes (mitophagy), iii) their replacement by mito-biogenesis [17]. In this highly regulated turnover, each step is required to successfully activate the next one; In particular, an effective mitophagy activates mitochondrial biogenesis, repopulating the pool of healthy mitochondria. To explain the presence of fragmented and exhausted mitochondria, we hypothesized alterations of mQC and mitophagy in FD cells. Indeed, we observed increased mitochondrial levels of DRP1 (**Fig. 2A),** alongside an increase of mitofusin (Mfn2) ubiquitination which tag fixed mitochondria for degradation (**Fig. 2B)** [18]. Accordingly, we detected an increase in FD mitochondrial lysates of both active and inactive forms of LC3 (I-II respectively), crucial regulators of auto(mito)phagosome formation (**Fig. 2C**). The concomitant accumulation of autophagosomes (**Fig. 2D**) strongly suggests that exhausted mitochondria are primed for degradation (early steps of mitophagy) but not effectively removed, with consequent accumulation of intermediate bodies. To confirm the impairment of mitophagic-flux in FD cells, we used mt-Keima (AAV-MitoKeima), a reporter that changes fluorescence (from green to red) when mitochondria are delivered to lysosomes [19]. In FD-Cells, the ratio of red (acidic) to green (neutral) fluorescence is reduced then WT cells, indicating that most mitochondria were not delivered to the lysosome for degradation **(Fig. S2 A-B)**. PGC-1a downregulation, mirrored by impaired mitochondrial biogenesis in FD MEFs (**Fig. 2E-F)**, confirmed that the incomplete mitophagy was unable to initiate mitochondrial renewal. Specifically, the downregulation of PGC-1a, master regulator of mitochondrial homeostasis [20, 21], may be a critical factor that further leads to mitochondrial function decline.

**Figure 1.**
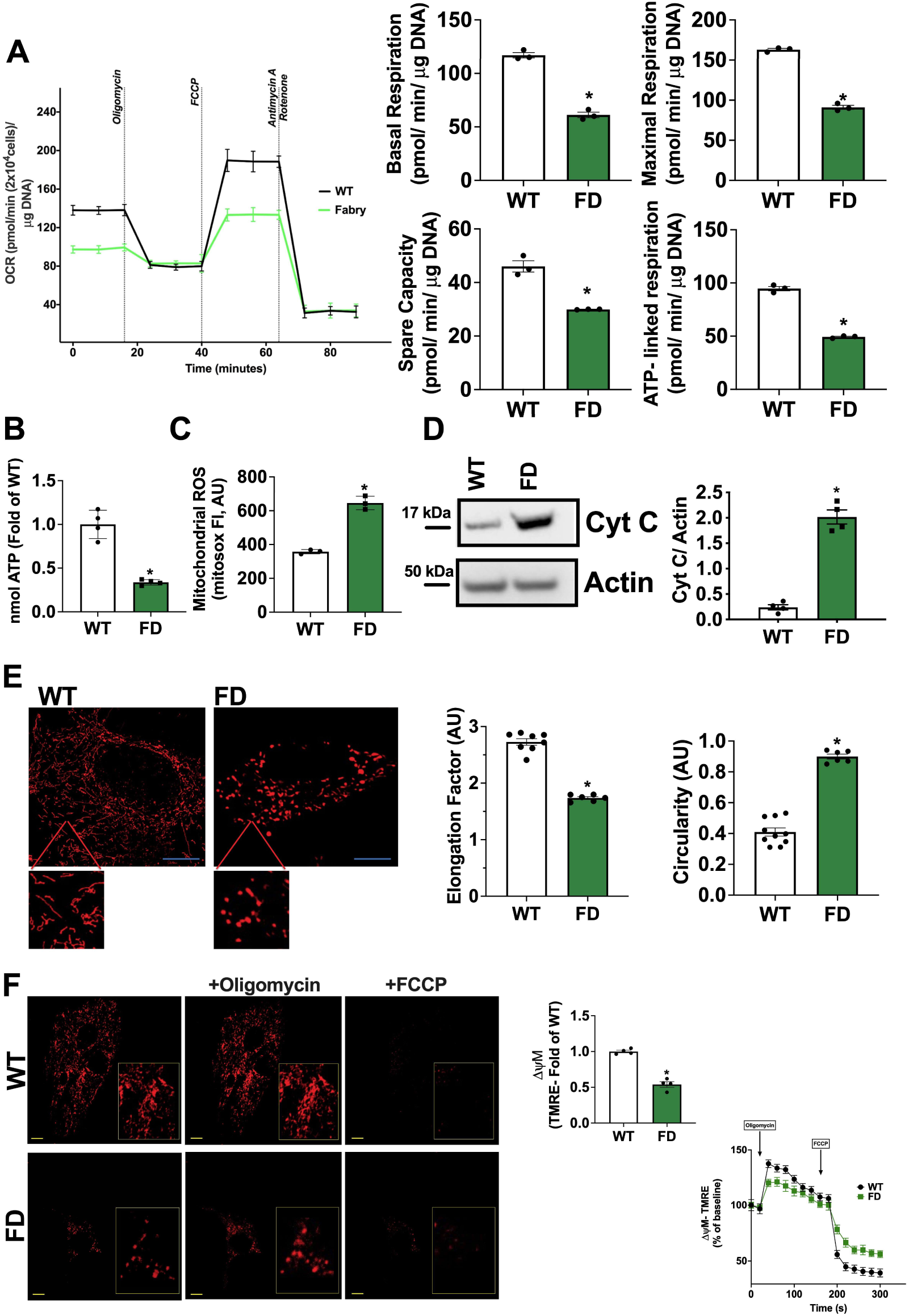
Alterations of mitochondrial function and dynamism in FD MEFs. Mouse Embryonic Fibroblasts (MEFs) were obtained from WT and FD mice. **(A)** The oxygen consumption rate (OCR) was assessed by Seahorse. Mitostress test displayed impaired mitochondrial performance in FD, with significant reduction of basal and maximal OCR, alongside spare capacity and ATP-linked respiration (WT n=3, FD n=3). **(B)** The levels of cellular ATP content were assessed by enzymatic assay and confirmed the energetic dysfunction in FD MEFs (WT n=4, FD n=4). **(C)** Mitochondrial ROS production was assessed in FD and WT-MEFs by using mitosox staining, and the results showed a significant ROS overproduction in FD-MEFs (n=3). **(D)** Western blot analysis of Cytochrome C on the cytosolic extract from WT and FD MEFs and relative quantification. Actin was used as loading control. Increased Cytochrome C release in the cytosol was observed for FD MEFs (WT n=4, FD n=4). **(E)** WT and FD MEFs were stained with mitotrackerred, and live cell imaging at high-resolution confocal microscope was performed to assess mitochondrial shape. Elongation factor and circularity were automatically calculated by microscope software (Nikon). In FD cells, mitochondria lost their elongated shape, and most mitochondria had high circularity, suggesting mitochondrial fragmentation (Elongation factor: WT n=8, FD n=6) (Circularity WT n=10, FD n=6) (Scale bar= 10 µm). **(F)** WT and FD MEFs were stained with TMRE to assess mitochondrial membrane potential and live cell imaging at confocal microscope was performed. Mitochondria are less polarized in FD cells (WT n=4, FD n=4). TMRE signal was also recorded in response to acute stimulation with oligomycin (5ug/ml) and FCCP (10uM) as control of dye direction (WT n=4, FD n=4) (Scale bar= 10 µm). Images are representative of at least 3 independent experiments; In the graph, Mean +/- SEM were plotted. Mann-Whitney U test was used to assess the significant differences between groups (*p<0.05 WT *vs*. FD).

**Figure 2.**
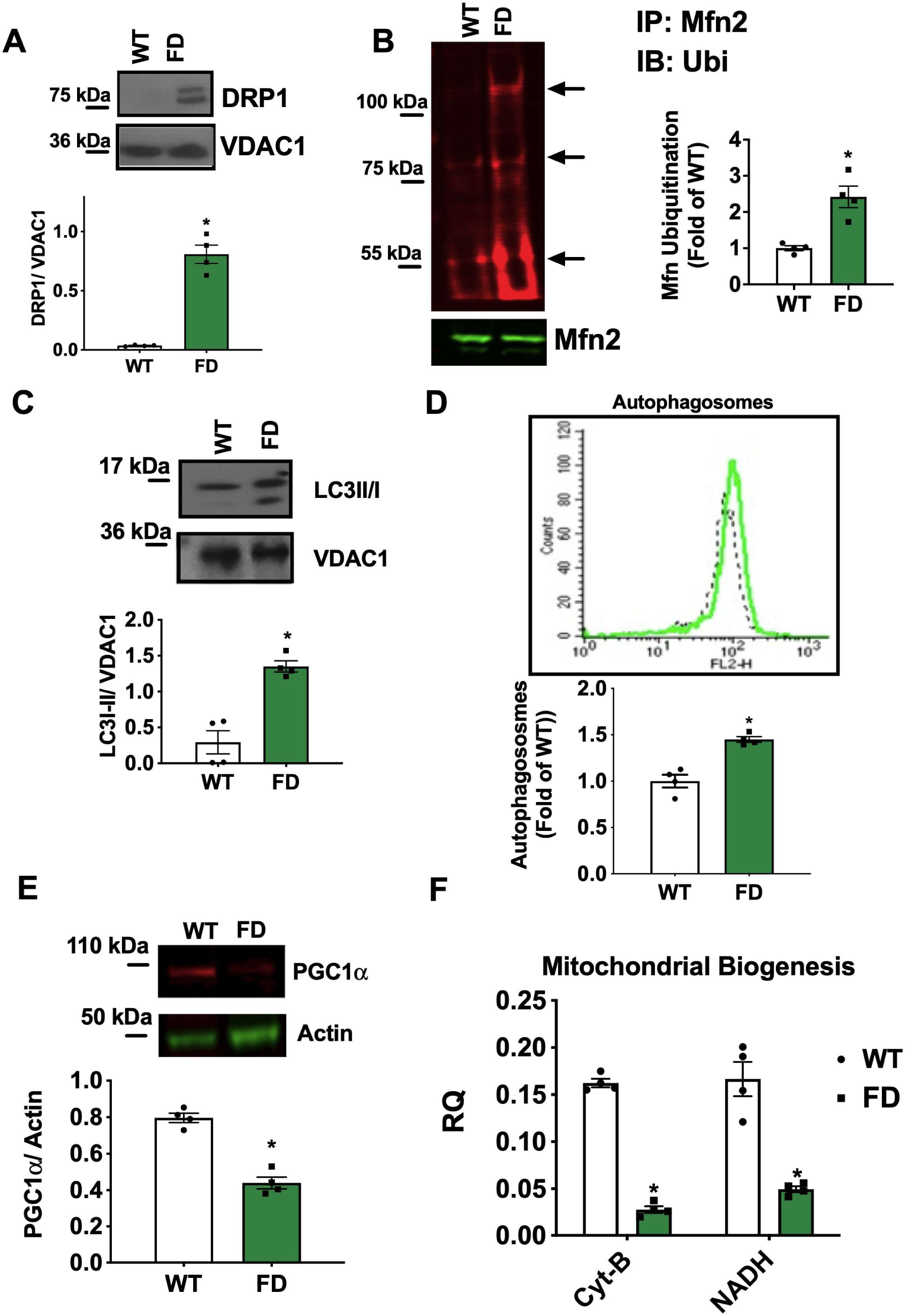
Alterations of mitochondrial turnover in FD MEFs. Mouse Embryonic Fibroblasts (MEFs) were obtained from WT and FD mice, and mitochondrial and whole cell lysate were performed to assess the levels of key markers of mitochondrial turnover. **(A)** Western blot analysis of DRP1 on the mitochondrial extract from WT and FD MEFs and relative quantification. VDAC1 was used as a loading control for mitochondrial preparation. Accumulation of DRP1 was detected in FD MEFs (WT n=4, FD n=4). **(B)** Ubiquitination test was performed to assess the levels of Mitofusin (Mfn) ubiquitination in WT and FD MEFs. Total immunoprecipitated Mfn levels were used as a loading control. Increased levels of ubiquitinated Mfn were detectable in FD-MEFs (WT n=4, FD n=4). **(C)** Western blot analysis of LC3II/I on the mitochondrial extract from WT and FD MEFs and relative quantification. VDAC1 was used as a loading control for mitochondrial preparation. On the mitochondrial surface of FD MEFs, an accumulation of LC3II/I was detectable (WT n=4, FD n=4). **(D)** FD and WT MEFs were stained with a fluorescent probe specific for Autophagososme detection. Cytofluorymetric analysis was performed to quantify the amount of autophagosomes. An accumulation of autophagosomes was detectable in FD-MEFs (WT n=4, FD n=4). **(E)** Western blot analysis of PGC-1α on whole cell extract of WT and FD MEFs and relative quantification. GAPDH was used as a loading control for mitochondrial preparation (WT n=4, FD n=4). **(F)** Real-time PCR was performed to assess the gene copies of mitochondrial NADH and Cit B, the index of mitochondrial biogenesis (CIT B WT n=4, FD n=4, NADH WT n=4, FD n=4). Reduced PGC-1α levels and relative mitochondrial biogenesis were detected in FD MEFs. Images are representative of at least 3 independent experiments; In the graph, Mean +/- SEM were plotted. Mann-Whitney U test was used to assess the significant differences between groups (*:p<0.05 WT *vs*. FD).

### Mitochondrial and Lysosome Mutual Interrelationship in FD cells

Mitochondria and lysosomes are functionally associated; two crucial points characterize their interdependency: i) lysosomes guarantee the scavenging of unhealthy mitochondria through mitophagy, ensuring the optimal rate of mitochondrial turnover. ii) mitochondria produce energy essential for ATP-dependent lysosomal acidification, facilitating the effective scavenging. Our findings reveal impaired mitophagy and a buildup of exhausted mitochondria in FD cells. We propose that energy deficits stemming from mitochondrial damage may worsen lysosomal dysfunction in FD cells.

To corroborate this hypothesis, we evaluated lysosomal pH in live cells by Lysotracker/ Lysosensor co-staining and we observed that the lysosome pH was significantly impaired in FD-MEFs (**Fig. S3A**) and the exposure to ATP was able to rapidly induce lysosomal acidification (**Fig. S3B**). Accordingly, when we treated MEFs with bafilomycin, a potent inhibitor of lysosomal vacuolartype H+-ATPase (v-ATPase), we observed a faster increase of lysosomal pH in FD-MEFs compared to WT cells, indicating a reduced activity rate of v-ATPase in FD cells (**Fig. S3C**).

To allow the conclusion that an intracellular energetic deficit contributes to lysosomal dysfunction, we show that when WT MEFs were pharmacologically deprived of mitochondrial ATP production by FCCP exposure, cellular ATP content was downregulated (**Fig. S3D**) and an increase of lysosomal pH occurred (**Fig. S3E**). This evidence links mitochondrial energetic defect and lysosomal dysfunction, supporting the theory that the energetic failure has an essential role as primary mechanism of lysosomal dysfunction in FD pathogenesis.

Collectively, the experimental data obtained in MEFs allow us to confirm the mechanistic hypothesis of mitochondrial suffering in FD: the initial lysosomal defect affects mitophagy, blocking organelle renewal, and compromising cellular energetics, which in turn, exacerbates lysosomal dysfunction, thus generating a vicious cycle of mito-lysosomal failure in FD-cells. We sought to verify this mechanism *in vivo*, in particular in adult heart and cardiomyocytes, known to have a high mitochondrial-based metabolism, and to determine the impact of this phenomenon on organ damage and cardiac manifestations in FD.

### Evaluation of cardiac phenotype of FD mice

To evaluate FD mice as a potential model for the hypothesis mentioned above, we first assessed the cardiac phenotype of our hR301Q Tg/KO (FD) mice by a life-course approach performing *invivo* and *ex-vivo* investigations at 3, 6, 9, and 12 months.

Echocardiographic analyses of the left ventricle (LV) did not reveal major morphological abnormalities or systolic dysfunction of FD mice compared with WT controls. Indeed, parameters of LV function such as fractional shortening (FS), and parameters of cardiac structure like left ventricular end-diastolic diameter (LVEDD), cardiac mass, and relative wall thickness (RWT) remained comparable between the two groups over time (**Fig. S4A–D**). However, beginning at 9 months of age, FD mice showed evidences of impaired diastolic function: the E/E’ ratio assessed by transmitral and tissue doppler analysis indicated reduced relaxation during the diastolic phase (**Fig. 3A**), while at 12 months of age, the experimental endpoint, a compensatory enlargement of the left atrium (LA) in FD mice (**Fig. 3B**) can be recorded. Interestingly, the mild interstitial fibrosis of the LV (**Fig. 3C**) is detectable starting from 9-months, and it is of larger evidence in the FD compared to the WT LVs.

**Figure 3.**
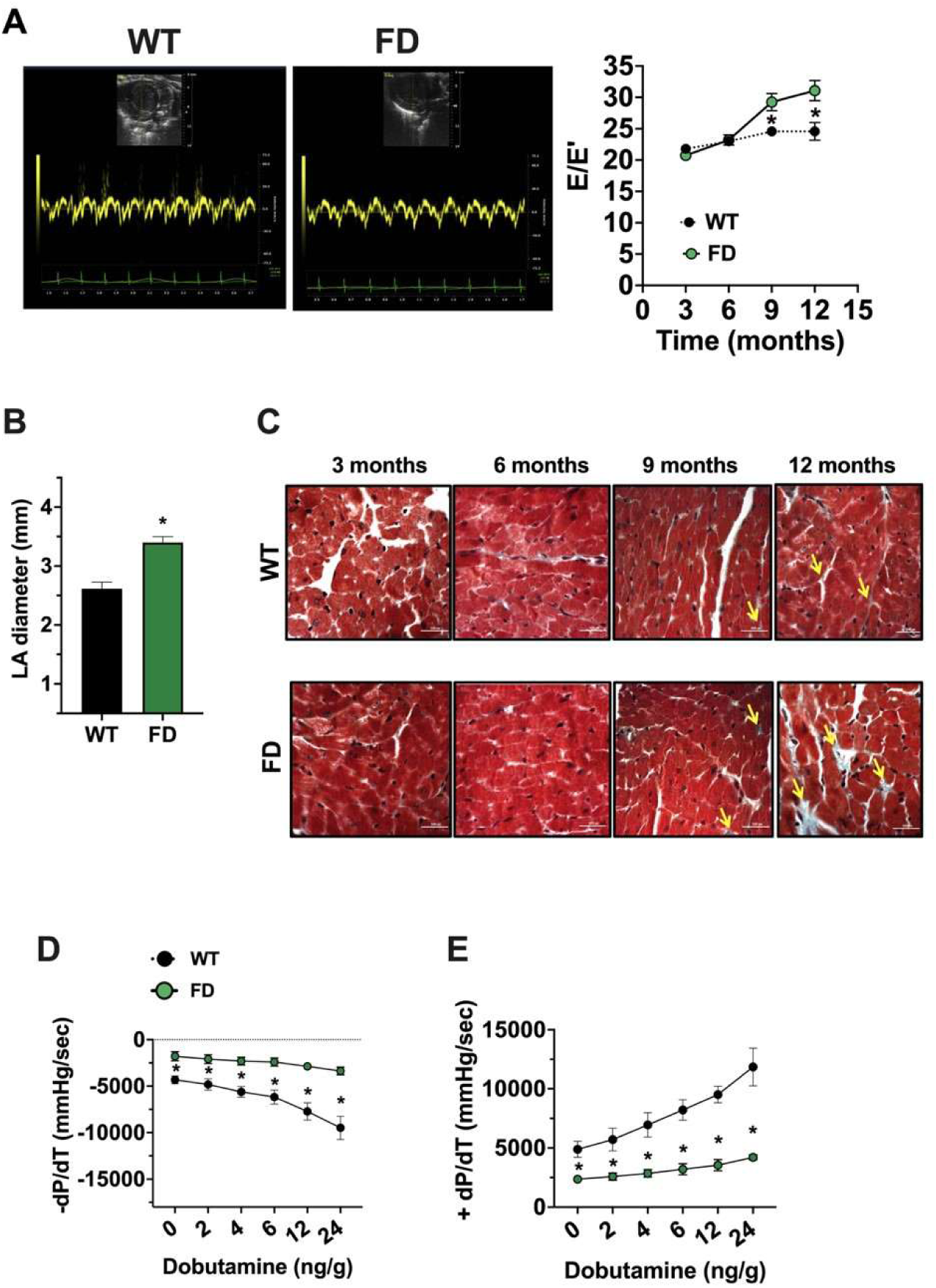
*In vivo* cardiac phenotype of FD mice Cardiac function was evaluated by cardiac ultrasounds in WT and FD mice of 3, 6, 9 and 12 months (WT n=16, FD n=18). (**A**) Analysis of transmitral flowmetry was performed, revealing that E/E’ ratio was increased in FD mice, index of diastolic dysfunction. (**B**) Antero-posterior diameter of Left Atrium (LA) was assessed at endpoint in WT and FD mice. An enlargement of LA occurred in 12 months old FD mice. Histological analysis was performed to assesses collagen deposition; **C**) Masson’s-trichromic staining evidenced increased fibrosis in cardiac tissue from FD mice starting from 9 months of age. Yellow arrows indicate interstitial fibrosis. Images were representative of at least 3 independent experiments (Scale bar= 50 µm). Hemodynamics responsiveness by invasive cardiac catheterization was performed in end point (12 months). **(D)** Left Ventricular (LV) dP/dt max and **(E)** LV dP/dt min in response to progressive sub chronotropic doses of IV Dobutamine were recorded in WT and FD mice and both indexes were impaired in FD mice (WT n=10, FD n=9). Data are shown as MEAN±SEM, WT (black), FD (green). *p< 0.05 FD *vs*. WT.

At 12 months we also assessed invasive hemodynamic responsiveness. **Table 1** summarizes the hemodynamic values of WT and FD mice at baseline. Interestingly, using this established technique [22], both the LV contractility (LV dP/dt max) and LV relaxation indexes (LV dP/dt min) were markedly impaired in FD mice in response to dobutamine challenge, compared to WT littermates (**Fig. 3D-E**). The intraventricular hemodynamics measurements were recorded using dobutamine at increasing doses that did not change HR throughout stimulation (**Fig. S4E**). Dobutamine-dependent increase of contractile and relaxation amplitude was significantly impaired in FD mice Overall, FD mice exhibited features consistent with diastolic dysfunction, including US tested left ventricular stiffness, left atrial enlargement, LV fibrosis and hemodynamic abnormalities, assessed by LV relaxation and contractility in response to dobutamine.

**Table 1.** Hemodynamic values of WT and FD mice at baseline. Experiments were performed in mice at 12 months of age. The data are reported as mean ± SD HR= heart rate LVSP= left ventricular systolic pressure; LVEDP= left ventricular end-diastolic pressure; LV dP/dt max = left ventricular maximal derivative of LV pressure; LV dP/dt min= left ventricular minimal derivative of LV pressure. *FD vs. WT p<0,02

|  | WT (n=8) | FD (n=7) |
| --- | --- | --- |
| HR (bpm) | 337±39 | 321±32 |
| LVSP (mmHg) | 99±11 | 73±6* |
| LVEDP (mmHg) | -2.5±4,6 | -2.5±3,6 |
| LV dP/dt max<br>(mmHg/s) | 4734±867 | 1927±502* |
| LV dP/dt min<br>(mmHg/s) | -4139±724 | -720±1419* |

### Progressive hypertrophy and contractile alterations of FD cardiomyocytes

Alongside fibrotic deposition, histological analysis of cardiac tissue revealed a progressive increase of FD cardiomyocytes (CMs) cross-sectional area in the intact FD cardiac tissue (**Fig. 4A**). Using a perfusion-based approach, we isolated adult ventricular CMs from FD and WT mice, in order to directly evaluate the morphology and the contractile performance of single cardiac myocytes. The morphometric analysis of isolated CMs confirmed the hypertrophic phenotype for FD-CMs starting from 9 months (**Fig. 4B**). Accordingly, we detected increased transcription of prohypertrophic genes, ANP and BNP, in line with the upregulation of MEF2, implicated in the gene expression program promoting cardiac hypertrophy (**Fig. 4C-D**). Hence, we evaluated the contractile performance of isolated CMs under field stimulation, both, in basal conditions and in response to adrenergic stimulation (isoproterenol). Data from single-cell contractility experiments indicate an increased basal contractility (**Fig. 4E**), and an impaired inotropic response to adrenergic stimulation of FD CMs (**Fig. 4F**) starting from 9 months of age.

**Figure 4.**
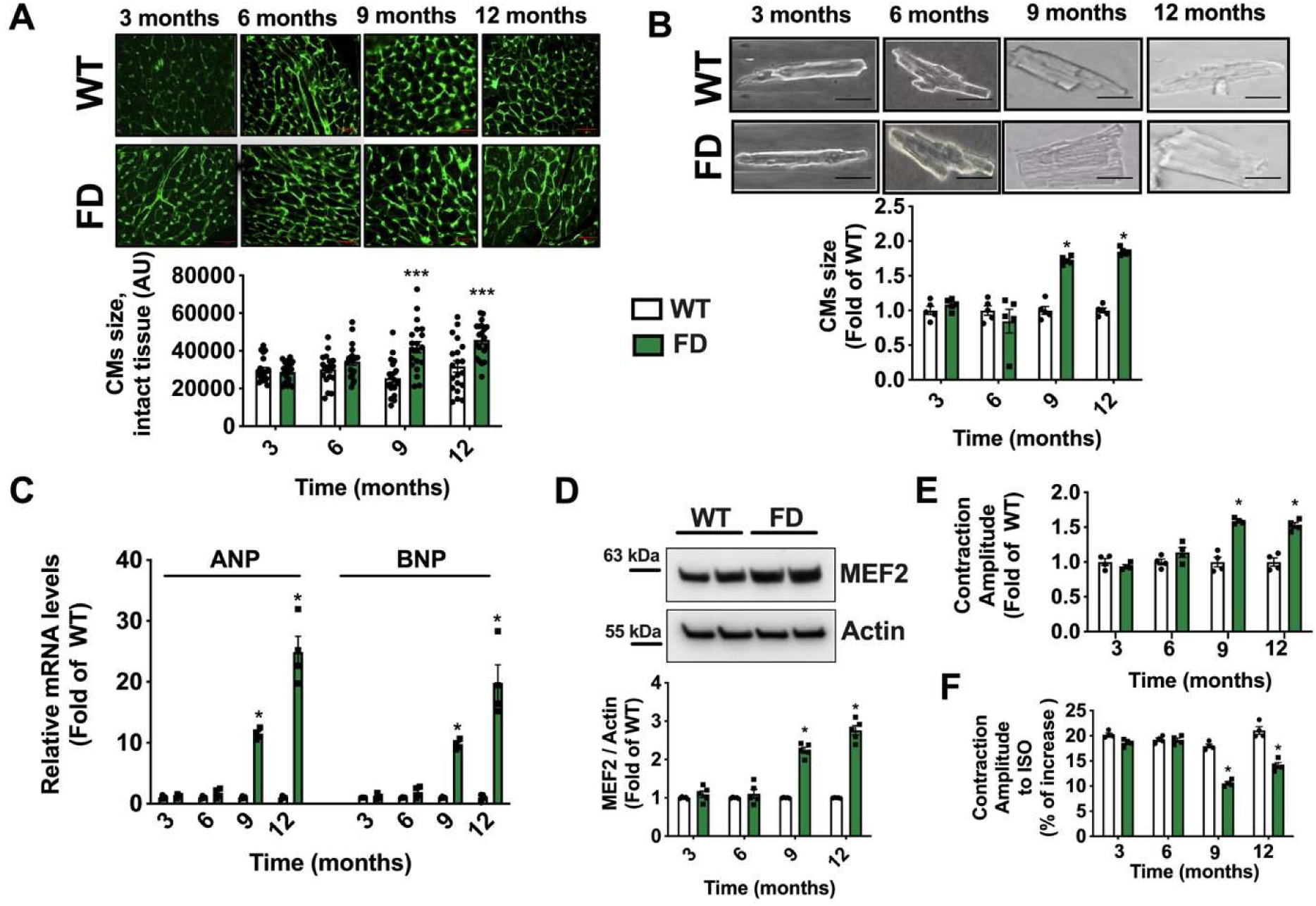
Hypertrophy and contractile abnormalities of FD cardiomyocytes. From WT and FD mice we isolated cardiac tissue and adult ventricular Cardiomyocytes to assess specifically cellular size and single-cell contractile profiles. **(A)** The WGA staining was used to evaluate the cross-sectional area of cardiomyocytes in intact tissue, showing the hypertrophic phenotype of FD cardiomyocytes starting from 9 months (WT n=20, FD n=20) (Scale bar= 50 µm). **(B)** Single cardiomyocytes (CMs) were isolated from adult WT and FD hearts, at different time points (3,6,9, and 12 months). The size of cardiomyocytes for each group was calculated and showed a significant increase in FD cell size from 9 months of age (WT n=5, FD n=5) (Scale bar= 20 µm). **(C)** Real-time PCR was performed to assess relative ANP and BNP mRNA levels in cardiac tissue from FD and WT mice. The transcription of both genes was significantly increased in FD heart starting from 9 months, according with the onset of CMs hypertrophy (WT n=4, FD n=4). **(D)** Western blot analysis was performed on WT and FD cardiac tissue to assess MEF2 protein levels, and the results displayed MEF2 upregulation in FD heart starting from 9 months (WT n=4, FD n=4). **(E)** Single-cell contractility analysis was performed under field stimulation. FD-CMs had a higher contractility than WT cells (WT n=4, FD n=4) probably attributable to their increased mass. **(F)** Stimulation with isoproterenol 100 nM was performed during the contractility recording. The results indicated an impaired responsiveness to adrenergic stimulation of FD CMs compared to WT cells (WT n=4, FD n=4). Images are representative of independent experiments; In the graph, Mean +/- SEM were plotted. Mann-Whitney U test was used to assess the significant differences between groups (*:p<0.05 WT *vs*. FD).

To note, in the context of isolated CMs the basal contractility can be only affected by cellular mass, thus explaining the different results recorded in whole mice, a complex model where systemic and pulmonary cardiovascular loads intervene (hemodynamics study) [23].

### Cardiac mitochondrial alterations precede the onset of myocardial dysfunction in FD mice

Our FD-mice model revealed that cardiac morphology and function, assessed *in vivo* and in isolated CMs, exhibited a delayed onset of cardiac symptoms starting at 9 months. To investigate the pathogenic role of mitochondrial alterations, we examined the mitochondrial phenotype in FD-hearts, focusing on the critical widow of 6-months age, which precedes the onset of clinical symptoms (9 months). Notably, mitochondrial energetics in FD-CMs were markedly compromised as early as 6 months, affecting basal and maximal respiration rates, spare capacity, and ATP-linked respiration (**Fig. 5A**), and increasing ECAR (**Fig. S1B**). To confirm the presence of an intrinsic mitochondrial defect in the Fabry heart, we performed high-resolution respirometry on freshly isolated cardiac mitochondria. Fabry mitochondria exhibited impaired State 2 (glutamate/malatedriven) and State 3 (ADP-stimulated) respiration, together with a reduced maximal electron transport system (ETS) capacity following FCCP uncoupling and a decreased coupling efficiency, as reflected by a lower respiratory control ratio (RCR) (**Fig. S5A**). Contextually, a downregulation of electron transport chain (ETC) complexes III-IV was detected (**Fig. S5 B**), corroborating the presence of dysfunctional mitochondria with impaired ATP production capability and abnormal ROS release in FD heart.

**Figure 5.**
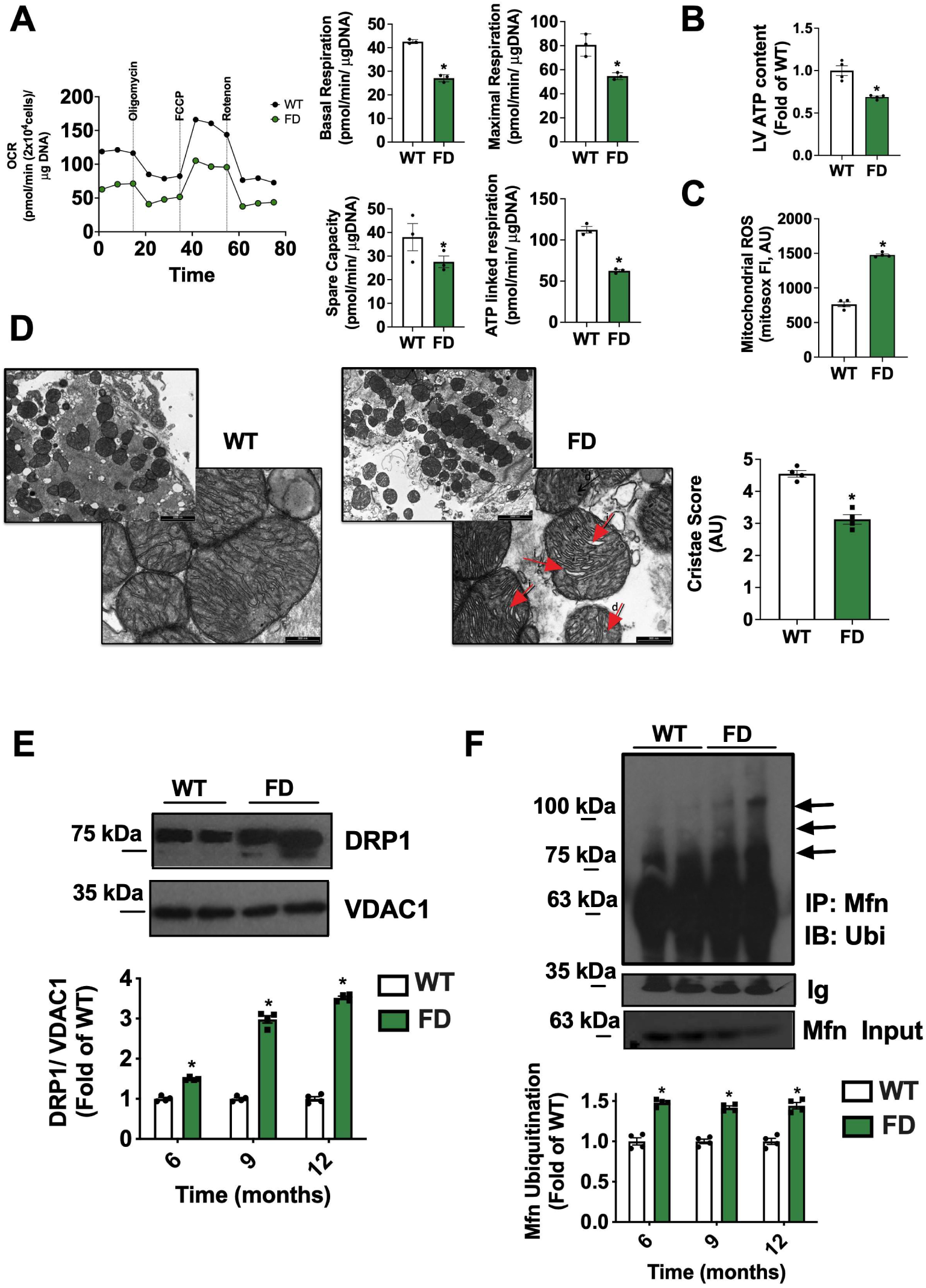
Alterations of mitochondrial function and dynamism in FD heart. From 6 months WT and FD mice we isolated adult ventricular Cardiomyocytes to assess the cardiac mitochondrial phenotype by reproducing key experiments performed in MEFs. **(A)** Oxygen consumption rate (OCR) was assessed by Seahorse, and mitostress test was performed to evaluate basal and maximal respiration rates, spare capacity and ATP-linked respiration. Mitochondrial performance was compromised in FD-CMs (WT n=4, FD n=4). **(B)** Total levels of ATP were determined in cardiac tissue from 6 months old WT and FD mice. A reduction of ATP reservoir was detectable in FD heart (WT n=4, FD n=4). **(C)** Mitochondrial ROS production was determined in isolated CMs by motosox staining and fluorescence was detected by cytofluorimetry (WT n=4, FD n=4). **(D)** Transmission electron microscopy (TEM) of left ventricle from 6-months-old WT and FD mice was performed to assess mitochondrial morphology and substructure. Representative images at two different magnifications revealed the accumulation of mitochondria with disarranged (d) and inflated (i) cristae (black arrows) (Upper panels, low magnification, scale bar= 2000 nm; bottom panels, high magnification, scale bar= 300 nm). Cristae score was calculated considering the percentage of mitochondrial area occupied by regular/ irregular cristaea, determined by imagej software application. Specifically, a Five-grade scoring system for cristae abundance and form with 0 as worst and 5 as the best was attributed (WT n=4, FD n=4). **(E)** Western blot analysis of DRP1 on cardiac mitochondria isolated from WT and FD mice from 6 months onward (6, 9, 12 months). VDAC1 was used as a loading control for mitochondrial preparation. Increased DRP1 levels were detectable in FD mitochondrial extract, for all time points (WT n=4, FD n=4). **(F)** The ubiquitination test revealed an increased Mitofusin (Mfn) ubiquitination in FD heart, confirming the alterations of mitochondrial dynamism (WT n=4, FD n=4). The blots are representative of the culminant changes in endpoint (12 months) while the quantitative analysis was shown for each time point. Images and graphs are representative of 4 independent experiments; Data in the graph are represented as Mean +/- SEM. Mann-Whitney U test was used to assess the significant differences between groups (*p<0.05 WT *vs*. FD).

Accordingly, the myocardial ATP content was significantly reduced in FD mice compared to controls (**Fig. 5B**), while a prominent mitochondrial ROS overproduction occurred (**Fig. 5C**).

We also detected abnormalities of cardiac FD mitochondria sub-structure, with disarranged and inflated cristae (**Fig. 5D**). In line with the data from FD-MEFs, mitochondrial levels of DRP1 and Mfn-ubiquitination were significantly increased in FD heart, thus confirming the accumulation of fragmented and exhausted mitochondria, targeted for degradation, in the heart as well (**Fig. 5E-F).**

We also observed the accumulation of LC3 in mitochondrial extracts, combined with the appearance of diffuse double-membrane autophagosome-like structures in FD myocardium (**Fig. S5 C-D**); contextually, we detected increased presence of autophagosomes in FD heart (**Fig. S5 E**), further supporting the impaired capacity of autophagosome scavenging, with accumulation of intermediate bodies. Moreover, PGC-1a levels were lower in the FD heart compared to WT, reflecting the decreased mitochondrial biogenesis (**Fig. S5 F-G**). It is noteworthy that all the described mitochondrial alterations occurred in FD heart starting not later than 6 months, so preceding the clinical manifestations (9 months). To better estimate when cardiac mitochondria get perturbed, we assessed mitochondrial performance also in neonatal and 2-months old CMs. Coherently with data form MEFs, early, in cardiac neonatal stage, we observed an impaired mitochondrial respiration and spare capacity, alongside mitochondrial ROS overproduction **(Fig. S6 A-B)**. However, the cellular ATP content was not affected, probably compensated by other mechanisms (i. e. higher glycolytic rate **(Fig. S1 C)**). The levels of PGC1-alpha starts to decline **(Fig. S6 C-D)**. In young CMs (from 2 months old mice) we observed a similar pattern, with the impairment of basal, maximal and ATP-linked OCR, and of mitochondrial spare capacity, alongside ROS overproduction **(Fig. S6 E-F)**, while energetic balance was not perturbed (ATP content) **(Fig. S6 G)**. PGC1alpha levels were constantly downregulated also in young FD hearts **(Fig. S6 H).** Cytochrome C release, a marker of leakage and terminal mitochondrial injury, occurred starting from 9 months (**Fig. S4 G**), according with the onset of organ damage. Hence, we evaluated if the release of Cytochrome C was activating CMs apoptosis; in FD hearts from 9 months old mice we detected a significant increase in cleaved Caspase 9 and 3, confirming the mitochondrial dependent (intrinsic) apoptosis activation **(Fig. S7 A-B).**

Overall, our results suggest an early and progressive mitochondrial impairment in FD heart which starts with reduced mitochondrial performance and increased susceptibility to stress (spare capacity) already in neonatal stage, and which proceeds impacting cardiac energetics (starting from 6 months), and cardiac homeostasis and function (9 months). Importantly, mitochondrial dysfunction persisted outside the cellular context, indicating a primary and direct mitochondrial abnormality that is independent of substrate availability and extramitochondrial regulatory pathways. Definitely, these data strongly support a pathogenic role of mitochondrial dysfunction in Fabry-associated cardiac manifestations.

### L-Arginine restores mitochondrial and energetic homeostasis in FD-MEFs

In our mechanistic hypothesis, unsuccessful mitophagy in FD cells not only induces the accumulation of unhealthy mitochondria but also prevents mitogenesis through PGC-1α downregulation, generating a mismatch between the cell metabolic demand and the ATP production alongside oxidative stress. A strategy that can comprehensively ameliorate mitochondrial homeostasis should be effective in preventing the energetic failure of FD cells, thus indirectly improving lysosomal function too. As such, L-Arginine (Arg) supplementation, able to induce PGC-1α upregulation, mitochondrial biogenesis and improvement of mitochondrial homeostasis [24–26], might prove the impaired energetic metabolism hypothesis true. Hence, we exposed FD-MEFs to Arg evaluating its effects on FD-related mitochondrial phenotype. In FD-MEFs treated with Arg, mitochondrial performance was significantly improved, in terms of basal and maximal respiration rate, spare capacity and ATP-linked respiration (**Fig. 6A**).

**Figure 6.**
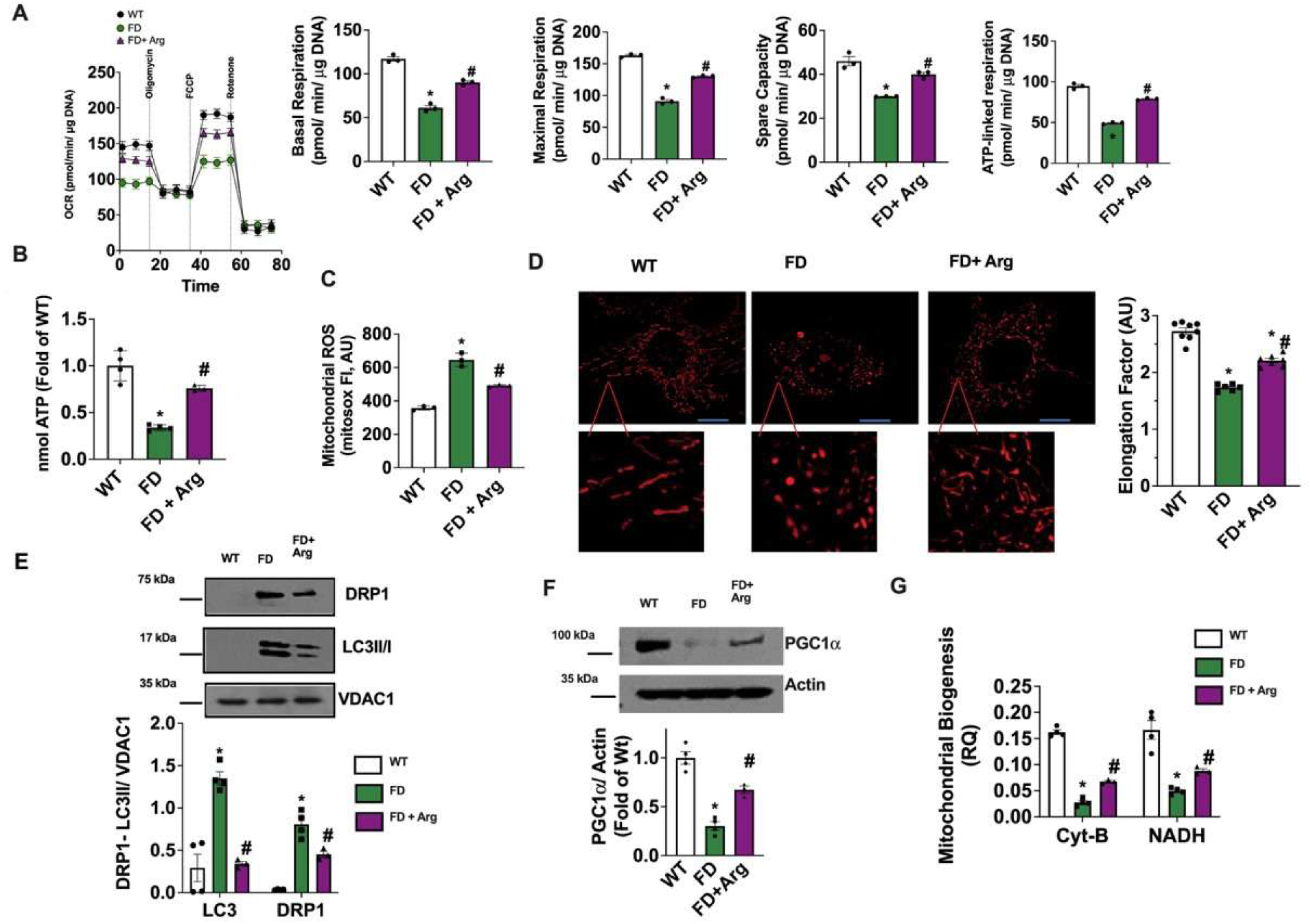
Arginine effects on mitochondrial homeostasis in FD-MEFs. FD and WT MEFs were exposed to 50 µM Arginine (Arg) for 24 hours and FD-related mitochondrial phenotype was assessed. (**A**) Oxygen consumption rate (OCR) was determined by Seahorse; mitostress test displayed a significant improvement of mitochondrial performance in response to Arg in FD-MEFs, with increase of basal and maximal OCR, spare capacity and ATP-linked respiration (WT n=3, FD n=3). (**B**) The total content of ATP was determined for WT and treated and untreated FD MEFs, and the results indicated that Arg exposure increased ATP reservoir (WT n=4, FD n=4, FD + Arg n=3). (**C**) Mitochondrial ROS production was assessed by mitosox staining and cytofluorimetric analysis in WT and FD-MEFs with and without Arg treatment. Arg exposure significantly reduced mitochondrial ROS production in FD-MEFs (WT n=3, FD n=3, FD + Arg n=3). (**D**) WT, treated and untreated FD-MEFs were stained with mitotracker-red, and live imaging at high-resolution confocal microscope was performed to assess mitochondrial shape. Elongation factor was automatically calculated by microscope software (Nikon). Arg treatment induced the recovery of functional mitochondrial elongated shape, with an increase of elongation factor (WT n=8, FD n=6, FD + Arg n=7) (Scale bar= 10 µm). (**E**) Western blot analysis of DRP1 and LC3II/I on mitochondrial extract from WT and treated and untreated FD MEFs, and relative quantification. VDAC1 was used as a loading control for mitochondrial preparation. Arg treatment induced the reduction of both DRP1 and LC3II/I mitochondrial levels, suggesting that it was able to counteract the alterations of mitochondrial turnover (LC3 WT n=4, FD n=4, FD + Arg n=3) (DRP1 WT n=4, FD n=4, FD + Arg n=3). (**F**) Western blot analysis of PGC-1a on whole cell lysate. Actin was used as the loading control. PGC-1a levels were recovered by Arg treatment (WT n=4, FD n=4, FD + Arg n=3). **(G)** Real-time PCR was used to assess the gene copies of mitochondrial NADH and Cit-B as an index of mitochondrial biogenesis. The results showed that Arg treatment improved mitochondrial biogenesis in FD MEFs (CIT-B WT n=4, FD n=4, FD + Arg n=3) (NADH WT n=4, FD n=4, FD + Arg n=3). Images are representative of at least 3 independent experiments; ANOVA followed by Bonferroni correction was used to assess the significative differences among groups. *p<0.05 WT *vs*. FD, # p<0.05 FD+Arg *vs*. FD.

This result was in line with Arg-induced recovery of mitochondrial membrane potential and of ATP cellular content (**Fig.S8A, Fig. 6B**) on one hand, and Arg-induced reduction of mitochondrial ROS levels on the other hand (**Fig. 6C**), suggesting that FD mitochondria started working properly again. The cellular antioxidant capacity, in terms of reduced vs oxidized glutathione ratio, was exhausted in FD-MEFs and restored in response to Arg treatment (**Fig. S8B**). Furthermore, Arg treatment counteract the accumulation of fragmented mitochondria (**Fig.6D**). The recovery of functional elongated shape of FD mitochondria was in line with the reactivation of the properly mitochondrial turnover suggested by reduction of DRP1 and LC3 mitochondrial levels (**Fig. 6E**), increase of PGC-1a expression and of mitochondrial biogenesis in FD-MEFs (**Fig. 6F-G**).

Concurrently, Arg supplementation was able to reduce autophagosome accumulation in FD-MEFs (**Fig. S9A**), as well as to improve lysosome acidification (**Fig. S9B**).

Overall, these results indicate that Arg widely restores mitochondrial homeostasis in FD cells guaranteeing a proper ATP production suitable for cellular processes, including lysosomal acidification and function. In turn, the recovery of lysosomal acidification and scavenging capacity can result in mitophagy unlock, saving the mitochondrial turnover.

### Arg supplementation prevents cardiac dysfunction in FD mice by promoting mitochondrial health

To evaluate whether Arg was able to prevent FD-cardiac manifestations, starting from 6 months of age, we supplemented FD mice with Arg. The treatment was able to prevent the onset of diastolic dysfunction in FD mice (**Fig. 7A**). The hemodynamic responsiveness of Arg-treated-FD mice was comparable to WT group; indeed, both systolic and diastolic indexes were preserved in FD mice supplemented with Arg (**Fig. 7B-C**). WGA staining showed a significant reduction of CMs size in the tissue from treated FD-mice (**Fig.7D**), and the ANP/BNP transcription was reduced (**Fig.7E**), overall suggesting that Arg treatment counteract the hypertrophic phenotype of FD-CMs. The contractile profile of CMs isolated from treated mice was preserved as well, in basal condition and in response to adrenergic stimulation (**Fig. 7F-G**).

**Figure 7.**
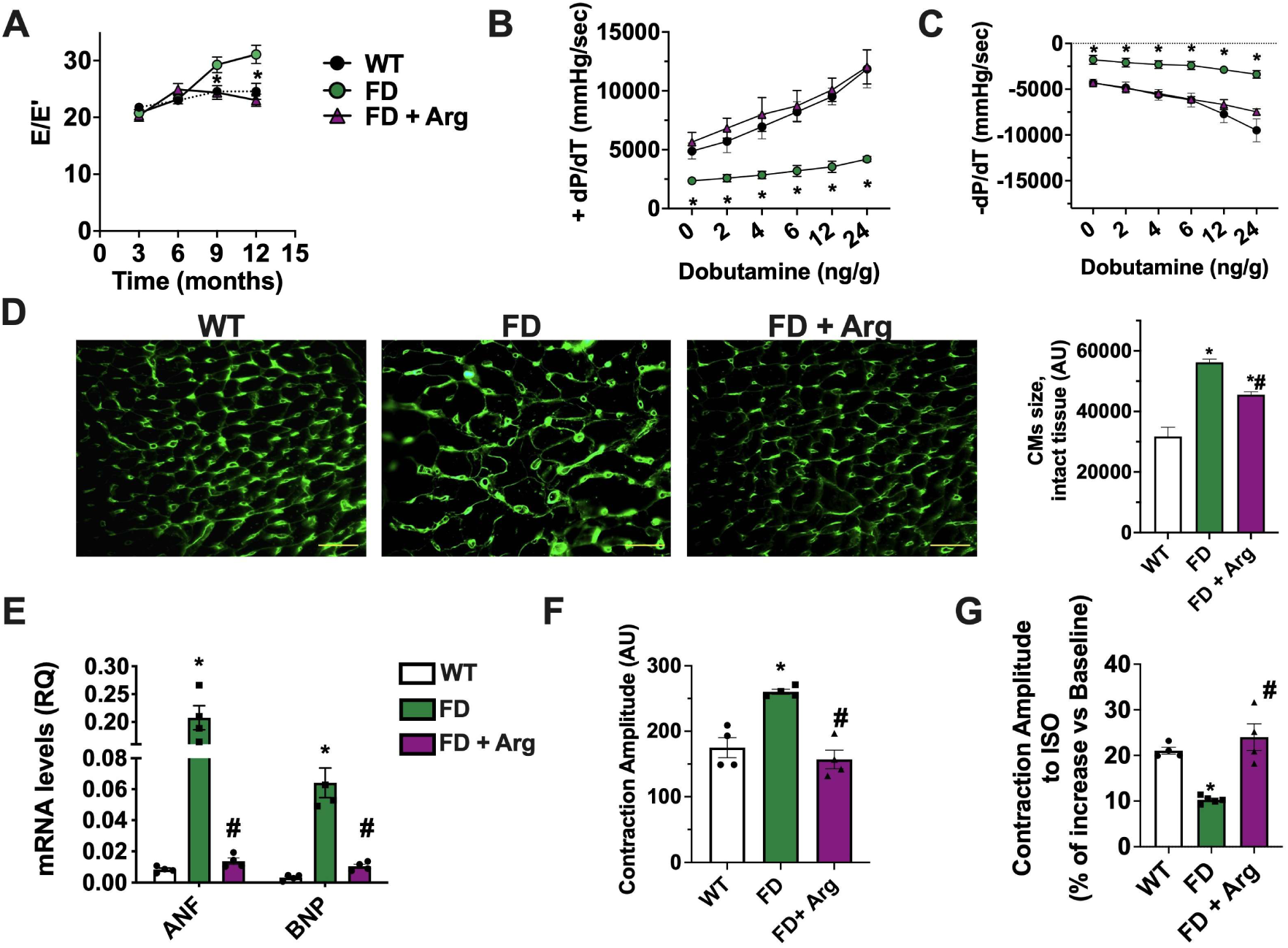
Arginine effects on cardiac phenotype of FD mice. Arginine (Arg) supplementation was tested in FD mice to evaluate its effects on cardiac phenotype. (**A**) A time course evaluation of Diastolic function was performed by transmitral flowmetry in FD-treated compared to untreated mice (WT n=13, FD n=15, FD+Arg n=10). We resumed data from Figure 3Efor WT and FD groups. Arg supplementation starts from 6 months of age, and here we reported also the echocardiographic data of the mice included in treated group at 3 and 6 months of age, before treatment initiation. The echo profile showed that the mice randomly assigned to the Arg-treated group, were indistinguishable from control mice at treatment baseline. The E/E’ ratio was increased in FD mice, but attenuated by Arg supplementation. (**B**) Left Ventricular (LV) dP/dt max and (**C**) LV dP/dt min in response to progressive doses of IV Dobutamine were recorded by cardiac catheterization in WT and FDtreated and untreated mice. Arg supplementation induced an improvement of both parameters in FD mice. For the WT and FD groups we re-proposed data from previous Figure 3F-G. (**D**) WGA staining on cardiac tissue section was perfomed to determine CMs size (Scale bar= 50 µm). The images quantification revealed that Arg treatment reduced cellular hypertophy in FD mice (WT n=5, FD n=5, FD+Arg N=5). (**E**) Real-time PCR was performed to assess ANP and BNP mRNA levels in cardiac tissue from WT and FD mice treated or not with Arg. Chronic Arg administration reduced ANP and BNP transcription levels in FD heart, in line with Arg induced effects on CMs hypertophy (WT n=4, FD n=4, FD+Arg N=4). (**F**) Adult ventricular CMs were isolated from WT and treated and untreated FD mice (WT n=4, FD n=4, FD+Arg N=4). Single-cell contractility analysis was performed under field stimulation. FD-CMs exhibited a higher contractility than WT cells while CMs from FDtreated mice had a normal basal contractility. **(G)** Stimulation with isoproterenol 100 nM was performed during the contractility recording. This analysis revealed impaired responsiveness of FD CMs, whilst the response to adrenergic stimulation is recovered in CMs from Arg-treated FD mice (WT n=4, FD n=5, FD+Arg n=4). Data are shown as mean±SEM. ANOVA followed by Bonferroni correction was used to assess the significative differences among groups. *p<0.05 WT *vs*. FD, # p<0,05 FD+Arg *vs*. FD.

The mitochondrial respiration rate was significantly improved in CMs isolated from Arg-treated FD-mice compared to the untreated group in terms of both basal and maximal respiration rate, spare capacity and ATP-linked-respiration (**Fig. 8A**). Accordingly, the content of myocardial ATP was preserved in treated FD-mice (**Fig. 8B**), while mitochondrial ROS were significantly reduced (**Fig. 8C**). In line with this prominent improvement of mitochondrial function, Arg exposure induced the recovery of cellular antioxidant capacity (**Fig. S8C**) and of ETC complexes III and IV expression in FD-myocardium (**Fig. S8D**).

**Figure 8.**
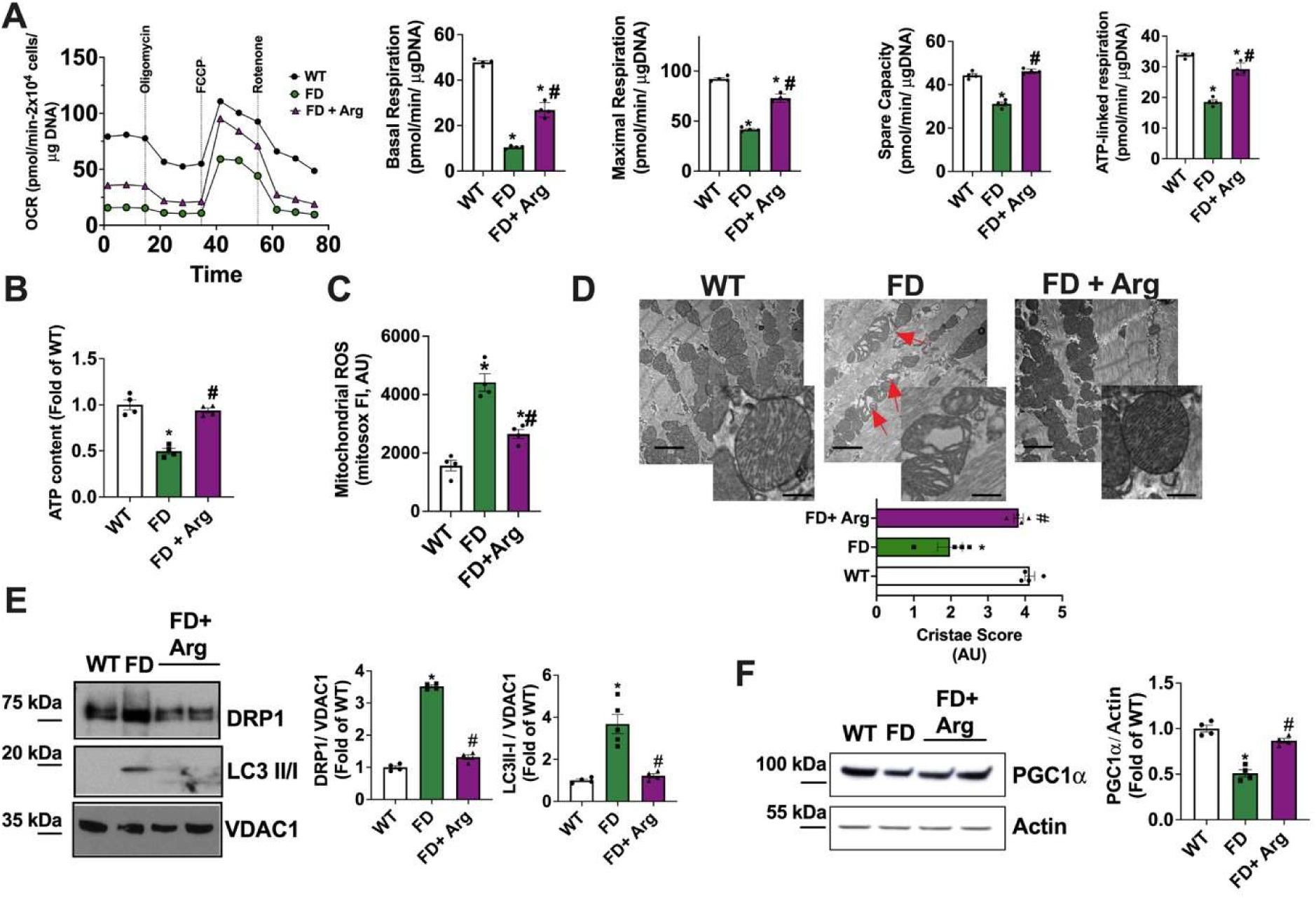
Arginine effects on the phenotype of cardiac mitochondria Heart tissue and CMs were isolated from WT, Arg-treated, and untreated FD mice to assess mitochondrial phenotype. (**A**) The oxygen consumption rate (OCR) was assessed by Seahorse. Mitostress test indicated that Arg supplementation improved basal and maximal mitochondrial respiration rate, spare capacity and ATP-linked respiration (WT n=4, FD n=5, FD+Arg n=4). (**B**) Total levels of ATP were determined in cardiac tissue from WT, Arg-treated, and untreated FD mice. Arg supplementation was able to increase the ATP content in FD hearts (WT n=4, FD n=5, FD+Arg n=4). **(C)** Mitochondrial ROS production was determined in isolated CMs by mitosox staining and cytofluorimetric analysis. The collected data indicated that Arg supplementation reduced mitochondrial ROS production in FD-CMs (WT n=4, FD n=4, FD+Arg n=4). (**D**) Transmission electron microscopy (TEM) of section of left ventricle from WT, Arg-treated, and untreated FD mice was performed to assess mitochondrial morphology and ultrastructure. Representative images at two magnifications (upper panels, low magnification, scale bar= 2000 nm; bottom panels, high magnification, scale bar= 300 nm), showed the presence in FD heart of mitochondria with disarranged cristae (Red arrows). Cristae score was calculated considering the percentage of mitochondrial area occupied by regular/ irregular cristae. Specifically, a Five-grade scoring system for cristae abundance and form with 0 as worst and 5 as the best was attributed and revealed that Arg supplementation recovered mitochondrial morphology (WT n=4, FD n=4, FD+Arg n=4). **(E)** Western blot analysis of DRP1 and LC3I/II in mitochondrial extract from WT, Arg treated and untreated FD hearts. VDAC1 was used as a loading control for mitochondrial preparation. Arg supplementation was able to reduce DRP1 and LC3II/I mitochondrial levels in the FD heart (DRP1 WT n=4, FD n=4, FD+Arg n=4) (LC3I/II WT n=4, FD n=4, FD+Arg n=4). (**F**) Western blot analysis was performed on cardiac mitochondria isolated from WT, Arg treated and untreated FD mice to assess PGC1a levels. Arg treatment induced PGC1a levels restoration (WT n=4, FD n=4, FD+Arg n=4). Images are representative of at least 3 independent experiments; ANOVA followed by Bonferroni correction was used to assess the significant differences among groups. *p<0.05 WT *vs*. FD, # p<0.05 FD+Arg *vs*. FD.

TEM analysis of cardiac tissue revealed that the physiological substructure of mitochondria was also recovered in Arg FD-mice (**Fig. 8D**). Ultimately, Arg supplementation was able to reduce mitochondrial levels of DRP1 and LC3, alongside the accumulation of autophagosomes (**Fig. 8E, Fig. S9C**). Moreover, PGC1a levels were increased in the heart of Arg treated mice (**Fig. 8F**). In line with the significant reduction of autophagosomes, Arg supplementation induced a prominent decrease of cardiac lysoGB3 accumulation (**Fig. S9D**), indicating that in turn, the amelioration of cardiac mitochondrial function can improve the lysosomal digestion properties, in accordance with data from MEFs.

### L-Arginine mediates mitochondrial rescue in FD by NO-dependent PGC1a upregulation

PGC1α critically regulates mitochondrial biology and ancillary programs relevant to mitochondrial homeostasis, including ATP production, ROS detoxification, and biogenesis [22].

Hence, we hypothesized that the pan-mitochondrial effects of Arg in FD are upstream mediated by its ability to induce PGC1α. Indeed, Arg is a potent NO precursor which in turn can increase PGC1α levels [22, 27–29] and support cardiac mitochondrial function [15]. To explore this possible mechanism, we first evaluated Arg mitochondrial effects in PGC1α-silenced FD cells, and our results showed that Arg did not improve mitochondrial function where PGC1α was not expressed (**Fig. S10 A-B**), confirming the hypothesis that Arg-action is PGC1α dependent. Hence, we verified the involvement of NO production in Arg-induced PGC1α mitochondrial improvement. First of all, we observed a reduced NO bioavailability in FD MEFs as well as in CMs, and Arg exposure was effective in inducing a significant increase in NO levels (**Fig. S11 A-B**). Arginine catabolism and ADMA levels were instead not different among WT and FD hearts (**Fig. S11 C**). When we blocked NO production in FD MEFs by chemical inhibition of NO-Synthase (NOS) activity, Arg was no longer able to upregulate PGC1α, as well as to improve mitochondrial respiration (**Fig.S11 D-E**). Opposite, NO donor mimics Arg effects in FD cells, similarly leading to PGC1α upregulation (**Fig.S12 A**). As a control, we also treated FD MEFs with citrulline, the other metabolite of NOS reaction, and we did not observe any PGC1α upregulation (**Fig.S12 B**). Previous reports indicate that the main biological effects of NO, including PGC1α upregulation, are mediated by direct cGMP-synthase activation and cGMP/PKG pathway evocation [22] [27] [28] [29]. To confirm that Arg induces PGC1α upregulation by activating NO/cGMP/PKG pathway, we used KT5823 to specifically inhibit PKG in Arg-exposed FD MEFs; as expected, Arg did not induce PGC1α upregulation when PKG was inhibited (**Fig. S12 C**). Opposite, when we treated FD MEFs with cGMP directly, PGC1α upregulation occurred, mimicking Arg action (**Fig. S12 D**).

It has been reported that Arg induces mTORC1 activation by acting on the lysosomal membrane protein Slc38a9 [30]. In turn, mTORC1 can regulate autophagy, lysosomal biogenesis as well as PGC1α activation and cellular metabolism [31]. Since mTOR dysregulation occurred in FD, we tested the possibility that the protective mitochondrial effects of Arg is mediated by mTOR signaling restoration in FD cells. Indeed, in FD MEFs we observed a reduced mTORC1 activity in terms of mTORC1 and S6K1 phosphorylation levels, and Arg exposure induced a significant increase of both, p-mTORC1 and p-S6K1 (**Fig. S13 A**). Accordingly, cardiac tissue from FD mice showed a significant reduction of p-S6K1 compared to wild-type tissue, while Arg restored p-S6K1 levels (**Fig. S13 B**). In both FD-MEFs and hearts, Slc38a9 protein levels remained unchanged in response to Arg (**Fig. S13 AB**).

When we inhibited mTORC1 with rapamycin in FD MEFs, we still observed Arg-induced PGC1α upregulation as well as improvement of mitochondrial respiration (**Fig. S13 C-D**).

## Discussion

The present study investigates the pathogenic role of mitochondrial dysfunction in FD. Specifically, we produced data in MEFs as well as in CMs isolated from a humanized mouse model of Fabry disease, that show altered mitochondrial turnover and accumulation of exhausted mitochondria, abnormal ROS production and cellular ATP collapse. Interestingly, mitochondrial failure strongly impairs the ATP-dependent lysosomal acidification and function, exacerbating in a vicious cycle, the primary lysosomal defect. In the heart, the mitochondrial alterations occur early (neonatal stage) and precede overt energetic failure (6 months) and later onset of cardiac phenotypes (9 months). This is the first study comprehensively assessing the cardiac phenotype of hR301Q Tg/KO mouse, which starting from 9 months of age, develops diastolic dysfunction, cardiomyocytes hypertrophy and contractile abnormalities. Noteworthy, the diastolic dysfunction [9, 32] in the absence of LV hypertrophy occurs in an early phase of cardiac involvement in FD patients, representing the right timing to identify early molecular targets and therapeutic interventions, corroborating the translational relevance of this mouse model. A similar phenotype of diastolic dysfunction was described also for GLA-KO mouse model [33]. Our data point to early mitophagy alterations that induce the accumulation of exhausted mitochondria and prevents the mitogenesis, mismatching the metabolic demand of the heart. Mitochondrial targeting by Arg supplementation restores PGC1-a dependent mitochondrial homeostasis reversing FD cardiomyopathy development.

To the best of our knowledge, the present work provides the first evidence that FD displays mitochondrial abnormalities, which are intimately linked to impaired lysosomal function and cardiac phenotypes.

Lysosomes are actively involved in mitophagy, which, in turn, stimulates de-novo mitochondrial biogenesis [13] [34]. In FD-cells we recorded an accumulation of ubiquitinated and fragmented mitochondria, alongside reduced autophagosome disposal and mitophagy flux. Contextually, we observed suppressed PGC-1α expression and impaired mitochondrial renewal, probably resulting from the auto/mitophagy perturbations. Our data in mice, are in line with the defective autophagosome maturation observed in cells derived from FD patients [35]. Consistently, also in human cells, decreased activity and expression of respiratory chain enzymes occur, alongside an altered ATP production and dysregulation of mitochondrial-related miRNAs [11] [10] [12] [36]. Here, we propose mitochondrial turnover alterations as fundamental mechanism of energetic and oxidative stress inducing cardiac damage in FD. Indeed, CMs hypertrophy could represent a compensatory response to inadequate energetic supply, derived from dysfunctional mitochondria. In addition, the oxyradical over-production from aberrant mitochondria can trigger activation of pro-hypertrophic genes [37, 38]. In this scenario, the pathogenic hypothesis of mechanical storage of Gb3 as the primary trigger of organ damage in FD appears to be extremely simplistic, especially for the FD heart, where the GB3 deposits represent only 1-2% of the total cardiac mass [9]. Indeed, cardiac damage in FD seems to be not sensitive to Gb3 targeting therapies [39, 40]. The long-standing knowledge gap between the GLA deficiency in the heart and the cascade of molecular events leading to cardiomyopathy, is the major obstacle for advancing therapeutic strategy to treat FD cardiomyopathy.

Our works challenges the dogma that substrate accumulation is the ultimate primary inducer of organ damage. Instead, we propose a novel energetic mechanism that clearly impacts CMs morphology and function. Specifically, using a life-long developmental approach in mice, we depicted the path of cardiac phenotype progression in relation to mitochondrial and energetic failure in FD: i) Perturbations of mitochondrial function and homeostasis are systemic and early, occurring in embryonic and neonatal cells (MEFs, neonatal CMs). ii) Impaired mitochondrial performance progressively impacts cellular energetics, reducing ATP availability (6 months) iii) The energetic collapse and the leakage of stressed mitochondria can induce apoptosis, maladaptive changes and functional decline of the heart (9 months), with CMs hypertrophy, contractile abnormalities and diastolic dysfunction (see **graphical abstract**).

It is well established that during development, the heart shifts from anaerobic metabolism in the fetal/neonatal stage to a highly oxidative, mitochondrial-dependent metabolism in the adult myocardium [41]. This shift tightens the coupling of oxidative phosphorylation and cardiac contractile function, increasing mitochondrial workload and ROS generation. Consequently, the adult heart is more susceptible to mitochondrial dysfunction, as confirmed by several genetic mitochondrial diseases that present with cardiac dysfunction in adult life [42–44]. In mammals, cardiac ATP content declines progressively with age [45]. An age-related decline in mitochondrial function, PGC-1α-mediated mitochondrial biogenesis, and mitochondrial quality control are the main underlying mechanisms [46].

Building on this, here we proposed a mechanistic model of pathogenesis and progression of FD cardiomyopathy; in the neonatal/young stage, the mitochondrial defect is present but does not yet impact cardiac energetics (less reliance on mitochondrial metabolism and compensation by other non-mitochondrial pathways). In the adult heart, the higher OXPHOS demand unmasks the dysfunction, inducing energetic failure and apoptosis. FD can be considered a phenotype of accelerated aging (mitochondrial decline, ATP decrease, cardiac cell loss). As in other mouse model of metabolic cardiomyopathies, the concomitant activation of cardiomyocytes hypertrophy and apoptotic pathway could explain the not significant increase in the overall heart size [47, 48]. Apoptosis and cell number reduction can involve mitochondrial ROS overproduction due to electron transport chain inefficiency, which damages mitochondrial and cellular components, thereby activating intrinsic apoptotic pathways [49]. This process reduces the number of viable cardiomyocytes while the remaining cells undergo hypertrophic growth as a compensatory response [50, 51].

In this scenario, mitochondrial targeting could be a promising approach for the treatment of cardiac injury in FD. In particular, we tested the hypothesis that acting upstream on PGC-1α recruitment, master regulator of mitochondrial biology, we could effectively induce mitochondrial homeostasis reprogramming [52]. Herein, we chose to explore Arg supplementation, which causes NO production and activation of PGC-1α, mitochondrial function, and cell metabolism [25] [53, 54]. Our data show that Arg restores mitochondrial health in FD-MEFs as well as in FD-CMs, inducing the rescue of cardiac phenotype. Specifically, here we demonstrate that Arg ameliorates mitochondrial homeostasis mainly by restoring PGC-1α levels, in a NO-dependent fashion. In particular, Arg increases NO availability and evocates NO/cGMP/PKG pathway which then culminates in PGC-1α upregulation as previously described [22, 27–29]. It has been extensively reported that cGMP and its primary effector PKG mediate the main biological actions of NO, with a central role in the maintenance of cardiovascular system [55]; indeed, enhancing cGMP-PKG signaling has emerged as a potent strategy to treat heart maladaptive remodeling and metabolic disarrangement, for the ability to mediate PGC-1α upregulation, mitochondrial ATP generation and oxidative stress reduction [29]. PGC-1α, in turn, regulates mitochondrial homeostasis and ancillary programs relevant to mitochondrial biology [52]. Similarly to our FD model, mice genetically lacking PGC-1α show poor contractile reserve and accelerated transition to failure during stress conditions [55]. Hence, our study, according with evidence coming from different contexts [17, 21], indicate that Arg is able to recovery the mitochondrial homeostasis by NO dependent activation of PGC-1α, thus reprogramming mitochondrial fate. Overall, the potent PGC1α-dependent effect of Arg also point out the key role of PGC1α downregulation in FD mitochondrial and energetic failure; mitophagy perturbation could be responsible for early PGC1α inactivation, which in turn, exacerbates the mitochondrial dysfunction and disarrangement [56], thus representing an hot target to save mitochondrial homeostasis in FD cells. Epigenetic events induced by mitochondrial stress, could be responsible of PGC1α downregulation in FD cells; however, the specific mechanisms remains to be elucidated.

The increased energetic availability in response to Arg, in turn improves lysosomal performance, supporting the ATP-dependent H+ pump function and lysosomal scavenging properties. Accordingly, our data show that Arg exposure can restore the lysosomal acidification in FD cells. This represents a critical point also for the current available therapies for FD treatment, including enzyme replacement therapies (ERTs) and chaperonin therapy. Indeed, abnormal lysosomal pH may interfere with ERT and chaperonin effectiveness in the lysosomes. Hence, an adjuvant intervention to restore lysosomal acidification could be crucial for amplifying ERT and other treatments efficiency. Given this unmet need, and supported by robust data, we propose Arg as a potential clinical intervention, also considering the non-supra-physiological dose tested in our study.

We cannot exclude that other pathways could be involved in Arg induced protective effect on cellular fate, including mTOR dependent signaling. However, here we focused on Arg direct mitochondrial effects, which seem to be independent from mTOR activation [30, 31, 57]. Our results on Arg effectiveness in the treatment of mitochondrial dysfunction in FD are consistent with the emerging evidence about its potent effect in supporting mitochondrial function and energetics in other pathological contexts, including in diabetic cardiomyopathy, and MELAS [53], with similar effect on ETC complex III expression and ATP production [53]. Notably, our group previously demonstrated that arginine (Arg), through the improvement of mitochondrial function, was able to counteract the diastolic dysfunction phenotype in db/db mice [15].

The NO deficit that we observed in FD MEFs as well as in CMs could explain the prominent beneficial effects of Arg on FD-related phenotypes. Accordingly, dysregulation of NOS-dependent NO production and NOS uncoupling have been extensively described in FD [58, 59]. Interestingly, similar NO deficiency occurs in mitochondrial disorders, and NO precursors like L-Arg, have been proposed as important treatment option for these conditions [60]. Oxidative stress and reduced NAD+ availability are among the main mechanisms responsible for NOS uncoupling and NO reduction in mitochondrial disorders [60]. We can speculate that also in FD similar mechanisms could be responsible for NO deficiency, as we did not observe Arg depletion or ADMA increase in our model. In addition to suggest Arg as novel potential treatment for FD cardiomyopathy, our data provide proof of the concept that mitochondrial targeting is effective in preventing FD cardiac involvement and supports the pathogenic role of mitochondrial alterations in FD cardiac injury. Our results open new research perspectives focused on the design and selection of novel drugs for mitochondrial and energetic targeting in FD. The implications of our study support the therapeutic targeting of cellular energy homeostasis to prevent and treat organ damage and failure in chronic diseases.

### Study limitations

We did not find frank left ventricular hypertrophy (LVH) in our FD mice. However, all the other signs of FD cardiomyopathy are clearly detectable, including cardiac Gb3 deposits, markers of cellular hypertrophy, diastolic dysfunction, and contractile issues. In humans, LVH typically occurs only in the late stage of cardiac damage, and in such cases, for instance in female, the loss of myocardial function is not necessarily associated with myocardial hypertrophy [61]. Innovative therapeutic interventions should be designed on early cardiac alterations, even to effectively prevent LVH. hR301Q Tg/KO mouse is particularly suitable for this purpose, allowing to identify the early pathogenic mechanisms that precedes the activation of the irreversible/maladaptive response of LVH.

## Materials and methods

A detailed section of materials and methods employed in the current study is available in supplementary material.

## Animal study

We employed hR301Q GLA Tg/KO as mouse model of Fabry disease (FD mice) mice, as previously acknowledged [14]. See Supplementary materials for details regarding this mouse model and control mice (WT). FD and WT mice were kept in a controlled environment, with 12:12 h light/dark period; 23°C; 55–60% humidity; and free access to water and chow. Our study examined male and female animals, and similar findings are reported for both sexes.

## Statistics

All values are presented as mean ± SEM or SD as specified in figure legends. Mann-Whitney Unpaired test was used as appropriated to compare differences between the two groups (WT vs FD). ANOVA was performed to compare the different parameters among the different groups. Bonferroni post hoc testing was performed as appropriate. A significance level of p < 0.05 was assumed for all statistical evaluations. Statistics were computed with GraphPad Prism software (Dotmatics, Boston, MA).

## Study approval

*In vivo* experiments were conformed to Animal Research Reporting of In Vivo Experiments (ARRIVE) guidelines, and approved by the Italian Ministry of Health, (n.158/2018-PR).

## Data availability statement

Relevant information about data available directly from the corresponding author

## Authorship contribution statement

JG: Writing original draft, Investigation, Data curation, Methodology, Conceptualization, AF, FAC, AB, RA, AV, ES, PC, VDA, NP, AP, SDA, FV, RP, ER, AB, Investigation, Data curation, Writing – review & editing. CP, AP, LS Data curation, Writing – review & editing. JS: Resources, Writing – review & editing. DS, GS, GI Writing – review & editing, Validation, Supervision, Project administration, Resources, Conceptualization.

## Founding and Acknowledgements

We thank Professor Avvedimento for his critical revision and helpful discussions. This work was supported in part by a collaborative research agreement between Amicus Therapeutics and GI. The authors thank Amicus Therapeutics for kindly providing us with the founders, and also thank Tobias Willer, PhD and Biliana Veleva-Rotse, PhD for advices on the experimental design and review of the manuscript. GI and JG was supported by Next Generation EU, National Recovery and Resilience Plan, Investment PE8 – Project Age-It: “Ageing Well in an Ageing Society”. DS is supported by FRA 54 2020. GS is supported in part by the National Institutes of Health (NIH): National Heart, Lung, and Blood Institute (NHLBI: R01-HL164772, R01-HL159062, R01-HL146691, T32-HL144456). The graphical abstract has been created in BioRender. avvisato, r. (2025) https://BioRender.com/czwmsvu.

## Declaration of interests

The authors have declared that no conflict of interest exists.

## References

1. Germain, D.P., et al., Use of a rare disease registry for establishing phenotypic classification of previously unassigned GLA variants: a consensus classification system by a multispecialty Fabry disease genotype-phenotype workgroup. J Med Genet, 2020. 57(8): p. 542–551.

2. Kubo, T., Fabry disease and its cardiac involvement. J Gen Fam Med, 2017. 18(5): p. 225–229.

3. Lin, H.Y., et al., High incidence of the cardiac variant of Fabry disease revealed by newborn screening in the Taiwan Chinese population. Circ Cardiovasc Genet, 2009. 2(5): p. 450–6.

4. Yim, J., et al., Fabry Cardiomyopathy: Current Practice and Future Directions. Cells, 2021. 10(6).

5. Messalli, G., et al., Role of cardiac MRI in evaluating patients with Anderson-Fabry disease: assessing cardiac effects of long-term enzyme replacement therapy. Radiol Med, 2012. 117(1): p. 19–28.

6. Sestito, S., et al., The heart in Anderson-Fabry disease. J Biol Regul, 2020. 34(4 Suppl. 2): p. 63–69.

7. Mehta, A., et al., Natural course of Fabry disease: changing pattern of causes of death in FOS - Fabry Outcome Survey. J Med Genet, 2009. 46(8): p. 548–52.

8. Yenercag, M. and U. Arslan, Tp-e interval and Tp-e/QT ratio and their association with left ventricular diastolic dysfunction in Fabry disease without left ventricular hypertrophy. J Electrocardiol, 2020. 59: p. 20–24.

9. Linhart, A. and P.M. Elliott, The heart in Anderson-Fabry disease and other lysosomal storage disorders. Heart, 2007. 93(4): p. 528–35.

10. Weissman, D., et al., Fabry Disease: Cardiac Implications and Molecular Mechanisms. Curr Heart Fail Rep, 2024. 21(2): p. 81–100.

11. Lucke, T., et al., Fabry disease: reduced activities of respiratory chain enzymes with decreased levels of energy-rich phosphates in fibroblasts. Mol Genet Metab, 2004. 82(1): p. 93–7.

12. Gambardella, J., et al., Mitochondrial microRNAs Are Dysregulated in Patients with Fabry Disease. J Pharmacol Exp Ther, 2023. 384(1): p. 72–78.

13. Audano, M., A. Schneider, and N. Mitro, Mitochondria, lysosomes, and dysfunction: their meaning in neurodegeneration. J Neurochem, 2018. 147(3): p. 291–309.

14. Gambardella, J., et al., Experimental evidence and clinical implications of Warburg effect in the skeletal muscle of Fabry disease. iScience, 2023. 26(3): p. 106074.

15. Fiordelisi, A., et al., L-Arginine supplementation as mitochondrial therapy in diabetic cardiomyopathy. Cardiovasc Diabetol, 2024. 23(1): p. 450.

16. Ikawa, M., N. Povalko, and Y. Koga, Arginine therapy in mitochondrial myopathy, encephalopathy, lactic acidosis, and stroke-like episodes. Curr Opin Clin Nutr Metab Care, 2020. 23(1): p. 17–22.

17. Suliman, H.B. and C.A. Piantadosi, Mitochondrial Quality Control as a Therapeutic Target. Pharmacol Rev, 2016. 68(1): p. 20–48.

18. Gegg, M.E., et al., Mitofusin 1 and mitofusin 2 are ubiquitinated in a PINK1/parkin-dependent manner upon induction of mitophagy. Hum Mol Genet, 2010. 19(24): p. 4861–70.

19. Guerra, F., et al., Synergistic Effect of Mitochondrial and Lysosomal Dysfunction in Parkinson’s Disease. Cells, 2019. 8(5).

20. Fernandez-Marcos, P.J. and J. Auwerx, Regulation of PGC-1alpha, a nodal regulator of mitochondrial biogenesis. Am J Clin Nutr, 2011. 93(4): p. 884S–90.

21. Patten, I.S. and Z. Arany, PGC-1 coactivators in the cardiovascular system. Trends Endocrinol Metab, 2012. 23(2): p. 90–7.

22. Zhu, G., et al., The mitochondrial regulator PGC1alpha is induced by cGMP-PKG signaling and mediates the protective effects of phosphodiesterase 5 inhibition in heart failure. FEBS Lett, 2022. 596(1): p. 17–28.

23. Garrett, A.S., et al., Isolated cardiac muscle contracting against a real-time model of systemic and pulmonary cardiovascular loads. Am J Physiol Heart Circ Physiol, 2023. 325(5): p. H1223–H1234.

24. Chen, X., et al., Arginine promotes porcine type I muscle fibres formation through improvement of mitochondrial biogenesis. Br J Nutr, 2020. 123(5): p. 499–507.

25. Gambardella, J., et al., Effects of Chronic Supplementation of L-Arginine on Physical Fitness in Water Polo Players. Oxid Med Cell Longev, 2021. 2021: p. 6684568.

26. Zhang, H., et al., Dietary Arginine Supplementation Improves Intestinal Mitochondrial Functions in Low-Birth-Weight Piglets but Not in Normal-Birth-Weight Piglets. Antioxidants (Basel), 2021. 10(12).

27. Bhargava, P., J. Janda, and R.G. Schnellmann, Elucidation of cGMP-dependent induction of mitochondrial biogenesis through PKG and p38 MAPK in the kidney. Am J Physiol Renal Physiol, 2020. 318(2): p. F322–F328.

28. Nisoli, E., et al., Mitochondrial biogenesis by NO yields functionally active mitochondria in mammals. Proc Natl Acad Sci U S A, 2004. 101(47): p. 16507–12.

29. Borniquel, S., et al., Nitric oxide regulates mitochondrial oxidative stress protection via the transcriptional coactivator PGC-1alpha. FASEB J, 2006. 20(11): p. 1889–91.

30. Wang, S., et al., Metabolism. Lysosomal amino acid transporter SLC38A9 signals arginine sufficiency to mTORC1. Science, 2015. 347(6218): p. 188–94.

31. Takahara, T., et al., Amino acid-dependent control of mTORC1 signaling: a variety of regulatory modes. J Biomed Sci, 2020. 27(1): p. 87.

32. Shanks, M., et al., Systolic and diastolic function assessment in fabry disease patients using speckle-tracking imaging and comparison with conventional echocardiographic measurements. J Am Soc Echocardiogr, 2013. 26(12): p. 1407–14.

33. Nguyen Dinh Cat, A., et al., Cardiomyopathy and response to enzyme replacement therapy in a male mouse model for Fabry disease. PLoS One, 2012. 7(5): p. e33743.

34. Yoo, S.M. and Y.K. Jung, A Molecular Approach to Mitophagy and Mitochondrial Dynamics. Mol Cells, 2018. 41(1): p. 18–26.

35. Chevrier, M., et al., Autophagosome maturation is impaired in Fabry disease. Autophagy, 2010. 6(5): p. 589–99.

36. Polillo, P.C. and L. Ferri, New Jersey standards affect dispensing of i.v. admixtures. Am J Hosp Pharm, 1987. 44(11): p. 2486–7.

37. Shah, A.K., et al., Oxidative Stress as A Mechanism for Functional Alterations in Cardiac Hypertrophy and Heart Failure. Antioxidants (Basel), 2021. 10(6).

38. Moustafa, A., et al., The MEF2A transcription factor interactome in cardiomyocytes. Cell Death Dis, 2023. 14(4): p. 240.

39. Keslova-Veselikova, J., et al., Replacement of alpha-galactosidase A in Fabry disease: effect on fibroblast cultures compared with biopsied tissues of treated patients. Virchows Arch, 2008. 452(6): p. 651–65.

40. Thurberg, B.L., et al., Cardiac microvascular pathology in Fabry disease: evaluation of endomyocardial biopsies before and after enzyme replacement therapy. Circulation, 2009. 119(19): p. 2561–7.

41. Ellen Kreipke, R., et al., Metabolic remodeling in early development and cardiomyocyte maturation. Semin Cell Dev Biol, 2016. 52: p. 84–92.

42. Sagar, S. and A.B. Gustafsson, Cardiovascular aging: the mitochondrial influence. J Cardiovasc Aging, 2023. 3(3).

43. Stoevesandt, D., et al., Cardiac manifestations in adult MELAS syndrome (mitochondrial encephalomyopathy with lactic acidosis and stroke-like episodes syndrome)-a cross-sectional study. Orphanet J Rare Dis, 2025. 20(1): p. 62.

44. Finsterer, J. and S. Kothari, Cardiac manifestations of primary mitochondrial disorders. Int J Cardiol, 2014. 177(3): p. 754–63.

45. Amorim, J.A., et al., Mitochondrial and metabolic dysfunction in ageing and age-related diseases. Nat Rev Endocrinol, 2022. 18(4): p. 243–258.

46. Chistiakov, D.A., et al., Mitochondrial aging and age-related dysfunction of mitochondria. Biomed Res Int, 2014. 2014: p. 238463.

47. Zhang, D., et al., Mitochondrial Cardiomyopathy Caused by Elevated Reactive Oxygen Species and Impaired Cardiomyocyte Proliferation. Circ Res, 2018. 122(1): p. 74–87.

48. Arany, Z., et al., Transcriptional coactivator PGC-1 alpha controls the energy state and contractile function of cardiac muscle. Cell Metab, 2005. 1(4): p. 259–71.

49. Hekimi, S., Y. Wang, and A. Noe, Mitochondrial ROS and the Effectors of the Intrinsic Apoptotic Pathway in Aging Cells: The Discerning Killers! Front Genet, 2016. 7: p. 161.

50. van Empel, V.P. and L.J. De Windt, Myocyte hypertrophy and apoptosis: a balancing act. Cardiovasc Res, 2004. 63(3): p. 487–99.

51. Toischer, K., et al., Differential cardiac remodeling in preload versus afterload. Circulation, 2010. 122(10): p. 993–1003.

52. Abu Shelbayeh, O., et al., PGC-1alpha Is a Master Regulator of Mitochondrial Lifecycle and ROS Stress Response. Antioxidants (Basel), 2023. 12(5).

53. Kim-Shapiro, D.B. and M.T. Gladwin, Arginine for mitochondrial oxidative enzymopathy. Blood, 2020. 136(12): p. 1376–1378.

54. Morris, C.R., et al., Impact of arginine therapy on mitochondrial function in children with sickle cell disease during vaso-occlusive pain. Blood, 2020. 136(12): p. 1402–1406.

55. Roszczyc-Owsiejczuk, K. and P. Zabielski, Sphingolipids as a Culprit of Mitochondrial Dysfunction in Insulin Resistance and Type 2 Diabetes. Front Endocrinol (Lausanne), 2021. 12: p. 635175.

56. Salazar, G., et al., SQSTM1/p62 and PPARGC1A/PGC-1alpha at the interface of autophagy and vascular senescence. Autophagy, 2020. 16(6): p. 1092–1110.

57. Summer, R., et al., Activation of the mTORC1/PGC-1 axis promotes mitochondrial biogenesis and induces cellular senescence in the lung epithelium. Am J Physiol Lung Cell Mol Physiol, 2019. 316(6): p. L1049–L1060.

58. Chimenti, C., et al., Increased oxidative stress contributes to cardiomyocyte dysfunction and death in patients with Fabry disease cardiomyopathy. Hum Pathol, 2015. 46(11): p. 1760–8.

59. Shen, J.S., et al., Tetrahydrobiopterin deficiency in the pathogenesis of Fabry disease. Hum Mol Genet, 2017. 26(6): p. 1182–1192.

60. Almannai, M. and A.W. El-Hattab, Nitric Oxide Deficiency in Mitochondrial Disorders: The Utility of Arginine and Citrulline. Front Mol Neurosci, 2021. 14: p. 682780.

61. Niemann, M., et al., Differences in Fabry cardiomyopathy between female and male patients: consequences for diagnostic assessment. JACC Cardiovasc Imaging, 2011. 4(6): p. 592–601.

